# Identification of Long Non-coding RNA Candidate Disease Genes Associated with Clinically Reported CNVs in Congenital Heart Disease

**DOI:** 10.1101/2024.09.30.615967

**Authors:** Jacqueline S. Penaloza, Blythe Moreland, Jeffrey B. Gaither, Benjamin J. Landis, Stephanie M. Ware, Kim L. McBride, Peter White, CCVM Consortium

**Affiliations:** The Office of Data Sciences, The Abigail Wexner Research Institute, Nationwide Children’s Hospital, Columbus, Ohio, United States of America; The Steve and Cindy Rasmussen Institute for Genomic Medicine, The Abigail Wexner Research Institute, Nationwide Children’s Hospital, Columbus, Ohio, United States of America; Department of Pediatrics, Indiana University School of Medicine, Indianapolis, Indiana, United States of America; Department of Medical and Molecular Genetics, Indiana University School of Medicine, Indianapolis, Indiana, United States of America; Department of Medical Genetics, Cumming School of Medicine, University of Calgary, Calgary, Canada; Department of Pediatrics, The Ohio State University College of Medicine, Columbus, Ohio, United States of America

## Abstract

**Background:** Copy Number Variants (**CNVs**) contribute to 3-10% of isolated Congenital Heart Disease (**CHD**) cases, but their roles in disease pathogenesis are often unclear. Traditionally, diagnostics have focused on protein-coding genes, overlooking the pathogenic potential of non-coding regions constituting 99% of the genome. Long non-coding RNAs (**lncRNAs**) are increasingly recognized for their roles in development and disease.

**Methods:** In this study, we systematically analyzed candidate lncRNAs overlapping with clinically validated CNVs in 1,363 CHD patients from the Cytogenomics of Cardiovascular Malformations (**CCVM**) Consortium. We identified heart-expressed lncRNAs, constructed a gene regulatory network using Weighted Gene Co-expression Network Analysis (**WGCNA**), and identified gene modules significantly associated with heart development. Functional enrichment analyses and network visualizations were conducted to elucidate the roles of these lncRNAs in cardiac development and disease. The code is stably archived at https://doi.org/10.5281/zenodo.13799847.

**Results:** We identified 18 lncRNA candidate genes within modules significantly correlated with heart tissue, highlighting their potential involvement in CHD pathogenesis. Notably, lncRNAs such as *lnc-STK32C-3, lnc-TBX20-1*, and *CRMA* demonstrated strong associations with known CHD genes. Strikingly, while only 7.6% of known CHD genes were impacted by a CNV, 68.8% of the CNVs contained a lncRNA expressed in the heart.

**Conclusions:** Our findings highlight the critical yet underexplored role of lncRNAs in the genomics of CHD. By investigating CNV-associated lncRNAs, this study paves the way for deeper insights into the genetic basis of CHD by incorporating non-coding genomic regions. The research underscores the need for advanced annotation techniques and broader genetic database inclusion to fully capture the potential of lncRNAs in disease mechanisms. Overall, this work emphasizes the importance of the non-coding genome as a pivotal factor in CHD pathogenesis, potentially uncovering novel contributors to disease risk.

## Introduction

Congenital Heart Disease (**CHD**) is one of the most common congenital malformations, with a birth prevalence of 1% worldwide.^1^ In 60-75% of CHD cases, the genetic cause of the disease is unknown.^2^ Copy Number Variants (**CNVs**), which are subchromosomal variations in the number of copies of DNA segments ranging in size from 1000 base pairs (**bp**) to several megabases (**Mb**), account for 3-10% of the pathogenic causes of isolated CHD.^2^ Some recurrent CNVs are associated with a specific type of CHD, including 22q11.2 Deletion Syndrome (also known as DiGeorge syndrome, a ∼3Mb microdeletion impacting *TBX1*, a T-box transcription factor required for normal development of the second heart field),^3^ Williams-Beuren Syndrome (deletion of 7q11.23 that leads to the haploinsufficiency of nearly 25 genes, including the *ELN* gene that codes for elastin, important for blood vessel elasticity and associated with aortic and pulmonary stenosis)^4^ and 1p36 Deletion Syndrome (the most common terminal deletion syndrome with cardiac findings that include congenital heart defects and cardiomyopathy)^5^. In addition to these clinical syndromes, CNVs are linked with a considerably greater risk of non-syndromic CHD.^6^ Nonetheless, many distinct CNVs are observed in patients who have the same CHD diagnosis, and often their contribution to CHD pathogenesis remains unclear. The precise mechanisms by which CNVs lead to non-syndromic CHD are still largely unknown, necessitating further investigation into their functional impact and the underlying biological pathways involved.

When considering the molecular genetics of CNVs in CHD, disease etiology can be attributed to changes in the number of gene copies within the CNV. While the established diagnostic approach in CHD primarily emphasizes well-known protein-coding genes, as evidenced by resources such as the CHDgene website (a curated list of 142 high-confidence protein-coding CHD genes),^7^ it is becoming increasingly apparent that variation in non-coding regions, which constitute about 99% of the genome, may contribute to the pathogenicity of CNVs.^8^ Approximately 60-80% of the human genome can be transcribed into non-coding RNAs (**ncRNAs**), including microRNAs (**miRNAs**), long non-coding RNAs (**lncRNAs**), small nucleolar RNAs (**snoRNAs**), and several other classes of RNA molecule that provide various regulatory and functional roles in the genome.

lncRNAs are RNA molecules longer than 200 nucleotides that usually do not have a coding sequence.^9–11^ They are involved in diverse regulatory roles in gene expression, such as modifying chromatin, regulating transcription and post-transcriptional processes, and regulating RNA processing.^9,10^ lncRNAs originate from various genomic regions, including intergenic, intronic, and overlapping regions with other genes.^11,12^ At the DNA level, lncRNAs influence chromatin architecture by recruiting chromatin-activating complexes or looping factors, thereby regulating target genes.^9^ They also play significant roles in alternative splicing by interacting with splicing factors to control transcription rates and ensure proper gene expression.^12^ Post-transcriptionally, lncRNAs interact with RNA-binding proteins through sequence or structural motifs to regulate mRNA splicing. Furthermore, lncRNAs can act as miRNA sponges, binding to miRNAs and preventing their interaction with mRNAs, which is crucial for gene expression.^9^ Lastly, lncRNAs regulate protein stability and degradation by altering their structure to interact with proteins involved in key signaling pathways. Together, these roles make lncRNAs vital for differentiation and development, impacting gene regulation at the DNA, RNA, and protein levels.

lncRNAs have been implicated in various biological processes and diseases. Their dysregulation can lead to a variety of conditions, including cardiovascular diseases,^13–15^ neurodegenerative disorders,^16–19^ and other complex diseases.^20–23^ While the role of lncRNAs in CHD is emerging,^13–15^ their function as key regulators of gene expression during heart development suggests that they could be critical players in the pathogenesis of CHD. The lncRNA *lnc-STK32C/RP11-432J24.5,* associated with responses to stress, stimuli, and the immune system, is differentially expressed in ischemic hearts.^14^ *CRMA* (also known as *CARMA* for *CARdiomyocyte Maturation-Assoicated lncRNA*), previously known as *MIR1-1HG-AS1*, is an antisense lncRNA that regulates NOTCH1 signaling by targeting the miRNAs *MIR1-1*, *MIR1-133a2*, and other ncRNAs.^24^ Notably, in vitro knockdown of CRMA promotes cardiac commitment, suggesting that its downregulation is crucial for directing progenitor cells toward a cardiac fate. The intergenic lncRNA *BANCR* (*BRAF-Activated Non-Protein Coding RNAs*; HSALNG0071734)^25^ has been reported to be upregulated in atherosclerotic plaques,^26^ and may influence the development of cardiomyopathy.^13^ This lncRNA encodes a short open reading frame that produces a microprotein that regulates various cellular functions, including contributing to heart muscle structural integrity and function. *BANCR* has been shown to facilitate cardiomyocyte migration and ventricular enlargement during heart formation in primates, with its expression confirmed to be higher during this critical period of heart development.^27,28^ ClinVar, a database of genetic variants, contains several reports of pathogenic CNVs impacting the 9q21.11-21.13 region, which includes the gene *BANCR*.^29,30^ However, these CNVs impact several genes in the region, and *BANCR* has not yet been directly associated with CHD.

Several CNVs impacting lncRNAs have been described in CHD patients. In DiGeorge syndrome, the 22q11.2 deletion spans the antisense lncRNA *lnc-TSSK2-8* (HSALNG0134007), which activates canonical Wnt signaling by protecting β-catenin from degradation.^31^ Previously, only *TBX1* was thought to be the disease-causing gene in 22q11.2, but evidence suggests that the loss of *lnc-TSSK2-8* contributes to the development of CHD.^32,33^ Another intergenic lncRNA associated with CHD is *lnc-NIPA1-4* (HSALNG0104472), located in the pathogenic CNV 15q11.2, where in vitro studies have shown that knockout of this gene significantly impacts cardiomyocyte differentiation.^34^

Despite their significance, the role of most lncRNAs in human health and disease remains largely unknown. This study aims to identify novel lncRNAs that may play a role in the pathogenesis of CHD by examining a large cohort of CHD patients with candidate lncRNAs within clinically validated CNVs (CNV-lncRNAs). By leveraging transcriptomic data from the developing heart and six other organs, we utilized a data mining method known as Weighted Gene Co-expression Network Analysis (**WGCNA**) to assess the expression patterns, interaction networks, and potential mechanisms by which CNV-lncRNAs may influence CHD. We identified several CNV-lncRNAs with strong associations with key developmental pathways implicated in CHD. Furthermore, to promote transparency and reproducibility, we have made the software code and methodologies used in this analysis publicly available (https://github.com/nch-igm/lncCHDNet/). By providing our codebase as a reusable resource, we aim to contribute to a more comprehensive understanding of lncRNA function in CHD and pave the way for future discoveries in the field.

## Methods

### Description and Structure of the CNV Dataset of CHD Patients

**Figure 1** gives an overview of the process we followed to identify lncRNAs impacted by CNVs that may play a role in the etiology of CHD. To elucidate the potential role of copy number changes in lncRNA genes in abnormal heart development, we leveraged the recently published CHD cohort from the Cytogenomics of Cardiovascular Malformations (**CCVM**) Consortium.^35^ This dataset contains 1,363 CHD patients with both an abnormal echocardiogram and an abnormal finding on chromosomal microarray analysis (**CMA**). CMA is a genome-wide technique that identifies genomic gains or losses (CNVs) and regions of homozygosity (**ROH**). It is recommended as a first-tier test for patients with neurodevelopmental disorders and congenital anomalies^36^ and has been integrated into routine practice at many pediatric cardiac centers for infants with severe CHD^37^. Utilizing a centralized echocardiography review, the CCVM registry meticulously categorized each patient’s CHD using a hierarchical classification to generalize the cardiac phenotypes.^35^ Specific structural heart defects identified through clinical evaluations included, but were not limited to, septal defects (**SDs**), conotruncal heart defects (**CTDs**), and left and right-sided obstructive lesions. All patients had clinically reported pathogenic CNVs as assessed by American College of Medical Genetics and Genomics (**ACMG**) guidelines^38^. Both syndromic and isolated cases of CHD are present in this cohort to allow for comprehensive phenotypic and genotypic correlations.

**Figure 1.**
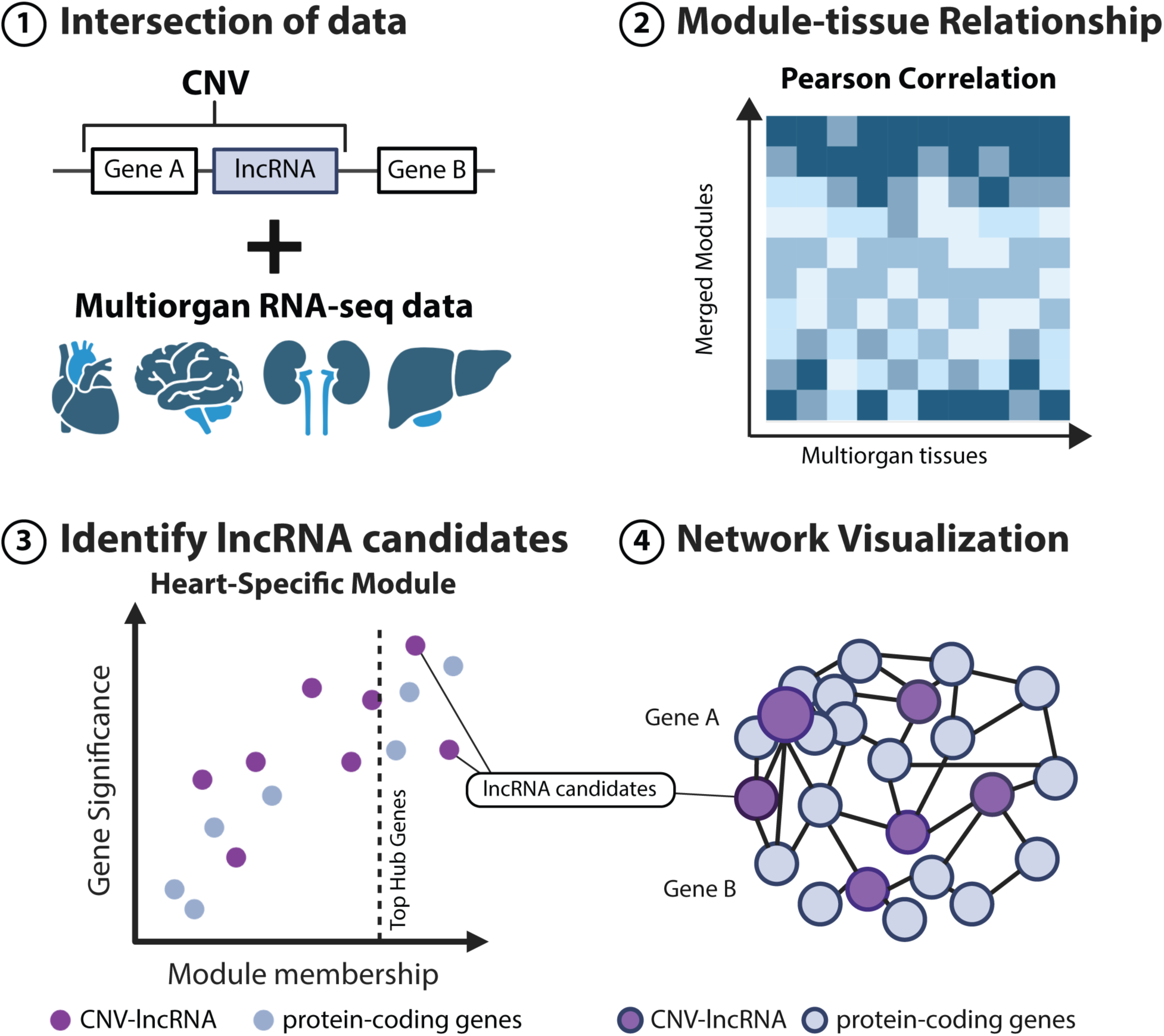
Workflow for Identifying lncRNA Candidates in CHD. This figure provides an overview of the multi-step methodological approach used in our analysis to investigate the roles of lncRNAs in CHD. In the first step, we integrate genomic information about lncRNAs from our cohort with expression data. In the second step, WGCNA is applied to identify modules that exhibit correlated expression across various tissues. The third step involves identifying hub genes, focusing on lncRNA candidates in the heart-specific modules. Finally, the network is visualized to highlight the interrelationships among CHD-related genes, lncRNAs, and protein-coding genes, showcasing the comprehensive methods applied in our analysis.

### Ethics approval and consent to participate

The CNV dataset applied in this study is available through the CCVM Consortium. The Consortium’s collection of CMA and clinical data was approved by each clinical center’s Institutional Review Board and utilized a waiver of informed consent. All data were anonymized before access and we followed general ethical guidelines for data use, including respecting privacy, data integrity, and intellectual property rights.

### Preprocessing of the CHD Cohort

The CCVM dataset underwent rigorous data wrangling and filtering to focus the analysis on isolated cases of CHD with CNVs. This filtering considered the CHD classification frequency, CNV length, location, and type (deletion, duplication, or ROH). The original dataset includes CMA data for 1,363 patients with CHD. Of these, 386 patients were classified as having syndromic CHD, while 977 were categorized as non-syndromic (**Supplementary Figure 1**). For this study, ‘syndromic’ refers to patients with genomic disorders with well-established associations with congenital heart defects, based on consensus opinions within the CCVM Consortium. The list of disorders considered syndromic was curated by the consortium and may not fully encompass all syndromic presentations. Across the non-syndromic patients, a total of 2,416 CNVs were recorded, including heterozygous (n=901) and homozygous (n=16) deletions, duplications (n=770), and runs of homozygosity (ROH, n=729) (**Supplementary Figure 2**). While most patients had a single CNV (n=963), 300 had two, and 100 had three or more CNVs listed on their clinical test report.

The first stage of preprocessing the CHD cohort was to remove all CNV entries capturing ROH. This type of CNV was excluded from our analysis, as discerning the function of a lncRNA contained within these regions would be considerably more complex than events that resulted in a loss or gain of a lncRNA. For the remaining 1,687 CNVs (duplications and deletions), the dataset was next filtered based on the size of the CNV to exclude small CNVs less than 5,000 bp and large CNVs greater than 5,000,000 bp (5 Mb). The lower limit for CNV length is determined by the size detection capabilities of the microarray technology. In contrast, the upper limit of 5 Mb is a commonly used cutoff to exclude aneuploidies and other large chromosomal rearrangements.^35^ The highly variable pseudoautosomal regions (PARs) on the X chromosome, which exhibit high rates of recombination and structural variation, were also removed to ensure more consistent and interpretable results in the analysis.

The final steps of preprocessing the data focused on the CHD category and defect classification. After removing ROH, small and large CNVs, and CNVs located in the PARs, 1,126 patients remained with 1,404 CNVs. Of these patients (821 non-syndromic; 305 syndromic), we removed all 305 syndromic patients and those with a diagnosis of cardiomyopathy (n=10) as this is primarily a disease of the heart muscle that may not directly relate to structural congenital defects. Patients with aortopathy (n=15) and arteriopathy (n=14) were removed due to the distinct pathophysiology of these conditions. Five other categories were removed due to their small sample size: CTD and atrioventricular septal defects (**CTD+AVSD**; n=15), Septal defect with Right Ventricular Outflow Tract Obstruction (**Septal RVOTO**; n=11), other (n=9), Patent Ductus Arteriosus (**PDA**; n=2), and Single Ventricle, other subtype (**SV,os**; n=2). The exclusion of these rare phenotypes enabled us to focus the analysis on the recurrent cardiac malformations that constitute the majority of phenotypes observed in this CHD cohort. The final dataset comprised 743 patients with 938 CNVs.

### RNA-seq Dataset and Processing for CNV-lncRNA analysis

We utilized the human organ RNA-seq time series of the development of seven major organs (ArrayExpress Accession Number E-MTAB-6814).^39^ The dataset covers 23 developmental stages across 7 organs (n=313: brain/forebrain 55, hindbrain/cerebellum 59, heart 50, kidney 40, liver 50, ovary 18, and testis 41), starting at four post-conception weeks (**PCW**) until 58-63 years of age. Normalized expression levels of lncRNAs calculated from this time series, expressed as transcripts per million (**TPM**), were obtained from the publicly accessible LncExpDB database (available at <u>lncRNA Expression Database</u>).^40^ For a description of how LncExpDB processed the transcriptomic data, see **Supplemental Methods**.

The RNA-seq data include gene expression levels for 101,293 lncRNAs and 19,957 protein-coding genes. When performing WGCNA, applying a minimum expression threshold of at least one transcript per million (**TPM**) is recommended, ensuring that only genes expressed in at least one sample are included in the analysis^34^. Applying this threshold to all samples in the developmental time series, 18,317 protein-coding genes had a minimum expression level of ≥1 TPM. For lncRNAs, we required a minimum expression level of ≥1 TPM in any of the 50 heart samples, as recommended for building the co-expression network. This filtering criterion identified 24,009 lncRNAs expressed in the heart at any developmental stage, along with corresponding expression levels for these genes across the other six organs.

### Identifying the CNV-lncRNA

Next, we wanted to identify all lncRNAs that fell within the 938 CNVs in the 743 isolated CHD patients in the processed CCVM cohort. Given the CNV coordinates of the original data were mapped to GRCh37, a liftover was performed to map regions to reference genome build GRCh38. The lncExpDB RNA-seq data were generated using a custom LncBook gene transfer format (**GTF**) file (LncBookv1.9_GENCODEv33_GRCh38.gtf). As such, this same file was used to identify CNV-lncRNAs. Although protein-coding genes are present in the GTF file, only lncRNAs were mapped to the CNV coordinates. Coordinates of CNVs were intersected with the LncBook lncRNAs utilizing the R package GenomicsRanges (v1.56.1). A total of 15,261 unique lncRNAs fell within clinically reported CNVs in the CCVM cohort. The median number of lncRNAs per CNV was 10, with an IQR of 24, and a maximum of 356 lncRNAs in a single CNV (**Supplementary Figure 3** shows the distribution of lncRNAs within the CNVs). Of the original 743 patients, 720 had a CNV overlapping 1 or more lncRNAs.

The list of 24,009 lncRNAs expressed in the heart at any developmental stage was intersected with the 15,262 unique lncRNAs falling within a CNV in the CCVM cohort. This resulted in a final list of 3,608 CNV-lncRNAs with evidence of heart expression, along with 18,317 unique protein-coding genes expressed in any tissue at any development stage, that together can be further analyzed in the co-expression network (**Figure 2E**). The median number of expressed lncRNAs per CNV was 2, with an IQR of 6, and a maximum of 110 expressed lncRNAs in a single CNV (**Supplementary Figure 4** shows the distribution of expressed lncRNAs within the CNVs). Of the original 743 patients, 557 had a CNV overlapping 1 or more expressed lncRNAs.

**Figure 2.**
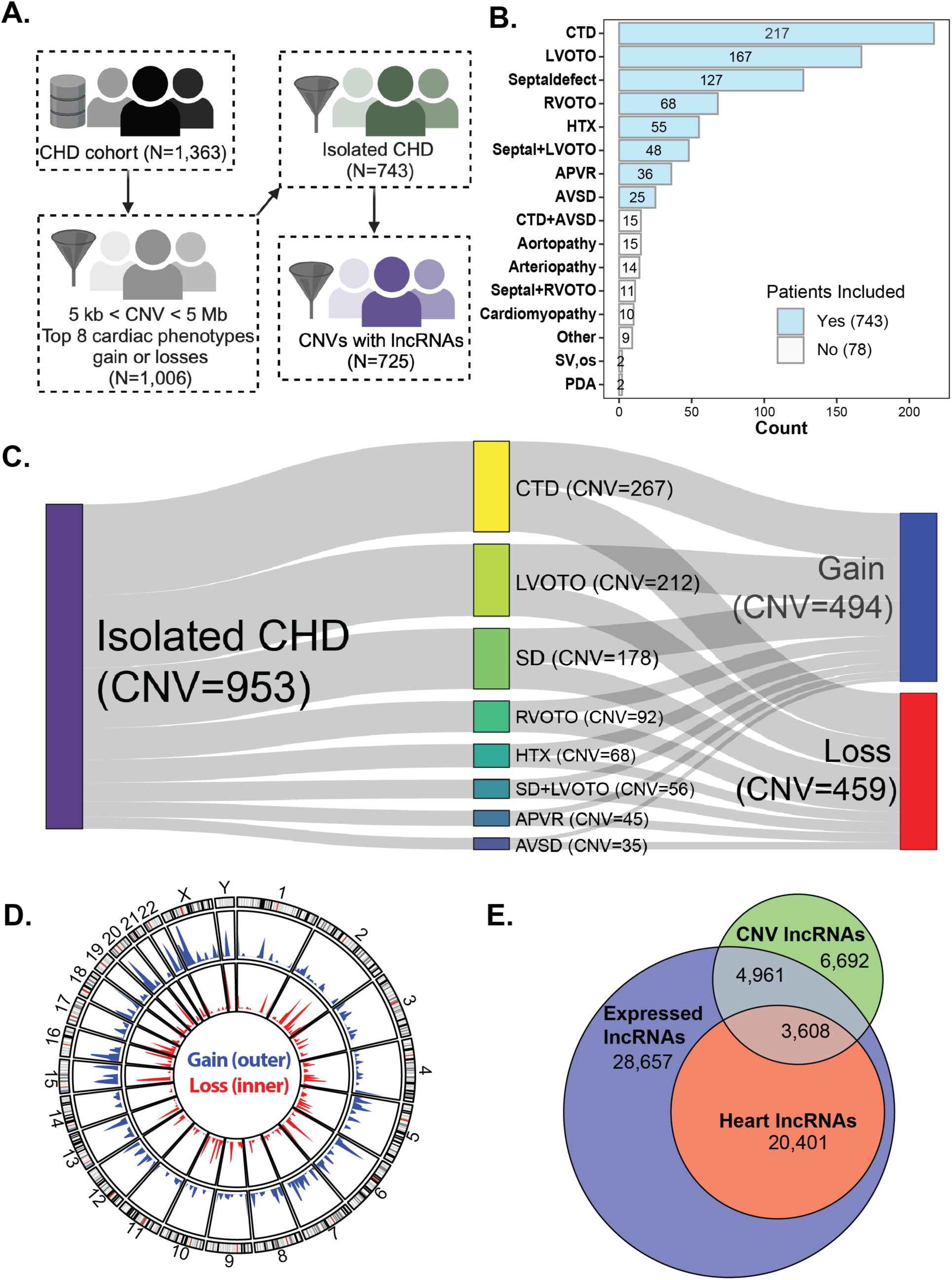
Overview of CHD Categories and CNVs in the CCVM Cohort. This figure details the selection of the CNV-LncRNAs reported in the CCVM cohort of isolated CHD patients. (A) Schematic representation of the CNV filtering process that refined the dataset for more targeted analysis. (B) The bar plot shows CHD category distribution in isolated (non-syndromic) CHD patients across the CCVM study cohort. Diagnoses are ranked by patient count, with bars indicating totals. Bars are color-coded: sky blue for patients included in the analysis and white for those excluded. (C) A Sankey diagram shows the distribution of isolated CHD cases in the cohort, detailing associated cardiac phenotypes and the total number of CNVs across all patients. Each node represents a category, diagnosis, or CNV type labeled with the total CNV count. The flow widths between nodes reflect the number of CNVs, highlighting the relative distribution of gains and losses across different CHD diagnoses in the cohort. (D) The circos plot visualizes CNV gains and losses across the human genome (hg38). The outer ideogram track displays chromosome bands, with black lines indicating gene-poor G-bands and red lines marking centromeres. The blue histogram track represents CNV gains, and the red track represents CNV losses. Taller bars highlight regions with higher frequencies of CNVs, pinpointing the genomic areas most impacted by gains and losses in the study cohort. (E) Venn diagram of expressed lncRNAs and CNV-associated lncRNAs illustrating the overlap between lncRNAs expressed in any tissue (blue), heart-expressed lncRNAs (red), and CNV-associated lncRNAs (green). The overlap regions indicate the number of lncRNAs shared between these categories, with the 3,608 heart-expressed lncRNAs used in our analysis being a subset of the total expressed lncRNAs.

### Network Construction

Protein-coding genes are well studied, and their functions are better understood than lncRNAs. Therefore, we can leverage the expression pattern of protein-coding genes to infer the function of lncRNAs. The final gene set consisted of 21,925 total genes, of which 18,317 are protein-coding genes expressed in any tissue, and 3,608 unique lncRNAs expressed in the human heart and falling within clinically reported CNVs in the CCVM cohort. We next constructed gene regulatory networks using lncRNAs and protein-coding genes identified through Weighted Gene Co-expression Network Analysis (**WGCNA**).^41^ WGCNA is a systems biology method that identifies clusters, or modules, of genes that show high levels of correlation in their expression patterns. These modules are then summarized using module eigengenes. A module eigengene captures the average expression profile of a gene module.^41^ This allows the module to be associated with different tissues, and hub genes specifically related to the heart to be identified. Building on this framework, we implemented stepwise module detection to refine our network analysis further, focusing on CNV-lncRNA co-expression networks to identify novel lncRNA hub genes.

In contrast to the block-wise module detection approach taken by Lu *et al.*,^34^ our study employs stepwise module detection for annotating CNV-lncRNAs. This approach offers greater control and precision in network construction by allowing the merging of similar modules. In contrast, the block-wise method does not and places genes into independent blocks, even when there may be potential similarities.^41^ For module detection, an appropriate soft-threshold power must be specified. WGCNA aims to create a network that follows a scale-free topology, characterized by a structure in which a few nodes are highly connected, and most nodes have low connectivity. We evaluated a range of powers, selecting a power value of 9 (R^2^=0.76), to ensure an optimal balance between achieving a scale-free topology and maintaining network connectivity (**Supplemental Figure 5**).

First, the topological overlap matrix (**TOM**) similarity was calculated with an unsigned network (see **Supplemental Methods** for details). The unsigned network considers positive and negative gene correlations but treats them equivalently without distinguishing their specific contribution.^41,42^ We chose an unsigned network to capture both activators and repressors of the heart developmental process. Second, network modules were detected by hierarchical clustering on the distance matrix derived from the TOM, specifically using 1 − *TOM* to measure gene-gene dissimilarity (**Supplemental Figure 6**), enabling genes to be grouped into co-expression modules based on their correlation patterns.^41^ Each module is assigned a unique identifier in the form of a color, which provides a visual and intuitive way to distinguish between different modules. These colors are arbitrary and serve as labels, but they remain consistent throughout the analysis, making it easy to track modules across different plots and analyses. For example, genes assigned to the “magenta” module exhibit similar expression patterns, while those in the “green” module form a distinct co-expression group. This color-based labeling system facilitates the interpretation of network topology and gene clustering results. Third, each module was summarized by its eigengene. Hierarchical clustering of these eigengenes, using dissimilarity measures (1 – correlation), was used to refine module granularity and merge similar modules at a 0.1 dissimilarity threshold (**Supplemental Figure 7**).

### Relate Modules to Organs and Developmental Stages

The final steps in our analysis focused on identifying which modules correlated most closely with the developing heart and identifying hub genes. First, the module-trait relationships were measured by calculating the Pearson correlation of the module eigengenes with the tissue type. To ensure robustness against Type I errors due to multiple comparisons across the seven organs, we applied a stringent p-value threshold of ≤ 0.001.^41^ Modules with high correlations to particular tissues, such as the heart, were highlighted as tissue-specific.

In addition to using the Pearson correlations, we visualized these relationships through a dendrogram, traditionally used to cluster co-expressed modules based on their eigengenes. In this case, we treated tissue types as an additional external trait and extended the dendrogram to reflect the similarity of module eigengenes to these tissue traits (**Supplemental Figure 8**). To complement the dendrogram, we also used an adjacency heatmap, which provides a visual representation of the correlation strengths between modules and tissues (**Supplemental Figure 9**). We focused on identifying modules most strongly correlated with the heart (**Supplemental Figures 10-12**), focusing on the 50 heart samples to developmental stages (**Supplemental Figures 13-15**).

An enrichment analysis was conducted on each WGCNA module to assess the representation of lncRNAs, protein-coding genes, and CHD genes. Hypergeometric and Fisher’s exact tests were employed to determine whether these gene types were significantly enriched or depleted within the modules. Fisher’s exact test p-values were adjusted for multiple testing, and the results were classified as “Enriched,” “Depleted,” or “Not Significant” (see **Supplemental Methods** for details.)

Next, we performed gene ontology (**GO**) and pathway enrichment analysis on the protein-coding genes within each module to identify overrepresented gene sets. This analysis focused on three GO categories: biological processes (**BP**), molecular functions (**MF**), and cellular components (**CC**), adjusting p values for multiple testing (see **Supplemental Methods** for details).

### Identification of Hub Genes

In WGCNA, a hub gene is a highly connected gene within a module, strongly correlated with the module eigengene, and often pivotal in the biological processes the module represents. Hub genes for modules of interest are identified by calculating the module memberships (**MM**) for all genes and selecting those with R^2^≥ 0.80.^43^ High MM values signify a strong and specific connection between a gene and a particular module. Low MM values indicate a weaker connection, showing that the gene’s expression pattern does not align closely with the module’s expression. In addition to MM, we calculated the correlation between each gene’s expression and the heart trait to determine gene-trait significance (**GS**) values. Here, we refer to the “heart trait” as a variable representing the expression data specific to heart samples across the dataset. GS represents how strongly each gene is associated with the heart trait, compared to other organs. Scatter plots were generated to visualize the relationship between MM and GS for each module, helping to identify hub genes with high MM and GS values. Gene aliases of the hub genes are identified by the LncExpDB database.^40^ The classification of lncRNAs among the hub genes was obtained from LncBook (Version 2.00).^44^ Additionally, LncExpDB was utilized to overview lncRNA tissue expression in organs not included in the developmental dataset.

To visualize the networks of correlated genes, the modules with significant association with the heart were exported to Cytoscape,^45^ where nodes represent genes and edges represent gene co-expression derived from the *TOM* matrix. We applied an edge weight minimum of 0.3 to refine the network, removing isolated nodes and small sub-networks for clarity. We focused on networks containing known CHD genes within the module of interest. To identify whether a protein-coding gene was a CHD gene, we cross-referenced our gene list with 142 high-confidence CHD genes from the CHDgene database. The CHDgene website (https://chdgene.victorchang.edu.au) is a valuable resource for clinicians and researchers, offering a curated list of genes with variants consistently shown to cause CHD in humans.

Embryonic heart Gini coefficient scores were used to quantify tissue-specific gene expression during human heart development (see **Supplemental Methods** for details).^28^ VanOudenhove *et al.* proposed that a higher Gini coefficient (e.g., >0.5) indicated a gene’s expression was more specific to the embryonic heart, identifying genes enriched for heart-related functions like TBX5, IRX4, and HAND1. To integrate this approach into the current analysis, LncBook identifiers were mapped to Ensembl gene accessions to retrieve Gini coefficients for genes with maximal expression in the embryonic heart.

## Results

The CCVM cohort comprises 1,363 patients and includes both syndromic and non-syndromic CHD (**Supplemental Figure 1**).^35^ We focused our analysis on patients with non-syndromic CHD, applying a filtering scheme that removed ROH, deletions less than 5 thousand base pairs (**Kb**), aneuploidies, and CNVs falling within the pseudoautosomal region (**Figure 2A**). The filtered dataset has 743 non-syndromic patients with a total of 938 CNVs larger than 5 Kb, but less than 5 million base pairs (**Mb**) (**Supplementary Figure 2**). Eight different subtypes of CHD were represented in the cohort: CTD, SD, left ventricular obstructive lesion (**LVOTO**), right ventricular obstructive lesion (**RVOTO**), SD + LVOTO, heterotaxy (**HTX**), atrioventricular septal defect (**AVSD**), and anomalous pulmonary venous return (**APVR**) (**Figure 2B**).

### Identification of CNV-associated lncRNA genes (CNV-lncRNAs)

The CCVM CMA data was successfully lifted from GRCh37 to GRCh38 to annotate lncRNA and protein-coding genes within each CNV using a comprehensive set of over 100,000 putative lncRNA genes from LncExpDB.^40^ Following the liftover process, some CNVs were split into multiple regions, resulting in a final dataset of 953 CNVs. The frequency of CNVs for the eight cardiac phenotypes is ranked from most to least in **Figure 2C**. There are 494 CNV gains (blue) and 459 losses (red). 573 patients had a single CNV, 140 had two CNVs, and 30 had three or more CNVs. The chromosomal distribution for gains and losses shows clinically reported CNVs in the CCVM non-syndromic CHD patients impact all chromosomes (**Figure 2D**). Overall, chromosome X had the most CNVs with 88 (60 gains and 28 losses). Chromosome Y had the least with 7 CNVs (5 gains and 2 losses). Only 72 of the 953 CNVs (7.6%) affected a known CHD gene. Of the 142 known CHD genes in the current CHDgene list, 33 (23%) were impacted by a CNV in our dataset. Notably, some of these known genes were recurrently affected, including *MYH11* (13 CNVs), *LZTR1* (8 CNVs), and *NOTCH1* (8 CNVs).

To identify CNV-associated lncRNAs (CNV-lncRNAs), we used RNA-seq data from a developmental time series of seven major human organs. For the WGCNA analysis, we first identified the subset of protein-coding genes expressed in any tissue across developmental stages. To focus on lncRNAs potentially involved in heart development and thus relevant to CHD, we restricted our analysis to lncRNAs expressed in the heart. We applied a minimum expression threshold of at least one TPM, resulting in 18,317 protein-coding genes and 57,627 lncRNAs that were expressed at any developmental time point across the seven different organs.

Among the expressed lncRNAs, 24,009 (42%) were expressed in the heart at some developmental stage. Of these heart-expressed lncRNAs, 3,608 (15%) were located within clinically reported CNVs from the CCVM cohort (**Figure 2E**). Out of the original 743 patients, 557 had CNVs overlapping one or more of these 3,608 heart-expressed lncRNAs, which we refer to as CNV-lncRNAs. These CNV-lncRNAs were used as input for the WGCNA analysis.

### WGCNA-Based Identification and Functional Analysis of Heart-Specific Gene Modules

For the WGCNA network construction, the TOM adjacency matrix contained a pairwise comparison for 21,925 total genes: 18,317 protein-coding genes expressed in any tissue and 3,608 CNV-lncRNAs derived from the intersection of lncRNAs expressed in the heart and CNVs in the CCVM cohort. The dendrogram constructed from hierarchical clustering of the distance TOM matrix contains information on the structure of gene expression correlations (**Supplementary Figure 6**). The hierarchical clustering constructs the modules for all genes. Each gene is tagged underneath by the color according to the module. For each module identified using the TOM-based clustering, eigengenes were calculated as the first principal component of the gene expression profiles, summarizing the overall expression pattern within each module. The hierarchical clustering initially resulted in 53 modules (**Supplementary Figure 7A**). Merging similar modules below the dissimilarity threshold of 0.1 led to 40 modules (**Supplementary Figure 7B**). All 40 modules and their associated counts of protein-coding genes, lncRNA genes, and known CHD genes are shown in **Supplemental Data Table S1**. Statistical analysis for gene enrichment of lncRNA genes and known CHD genes was performed using the hypergeometric distribution. Six modules (grey, magenta, paleturquoise, green, lightcyan1, and lightyellow) were enriched for lncRNAs. Four modules demonstrated enrichment of known CHD genes (darkturquoise, darkolivegreen, purple, and magenta). The magenta module (ME03) was the only module to display significant enrichment for both lncRNAs (p = 8.09 x 10^-08^) and known CHD genes (p=0.0112).

Pearson correlation values for the module eigengene-organ association across the 7 tissues for the 40 modules are depicted in the heatmap in **Figure 3** (modules and their associated correlations and p-values are available in **Supplemental Data Table S2**). The stronger the correlation, the higher the intensity of the color, with red representing positive correlations, blue representing negative correlations, and white representing no correlation. Modules with high positive or negative correlations (|r| > 0.6) and p-values below 0.001 are considered tissue-specific. For example, module ME40 (pink) is highly specific to the liver (r = 0.96, p = 5 x 10^-174^), while module ME28 (royalblue) is highly specific to the kidney (r = 0.83, p = 3 x 10^-82^).

**Figure 3.**
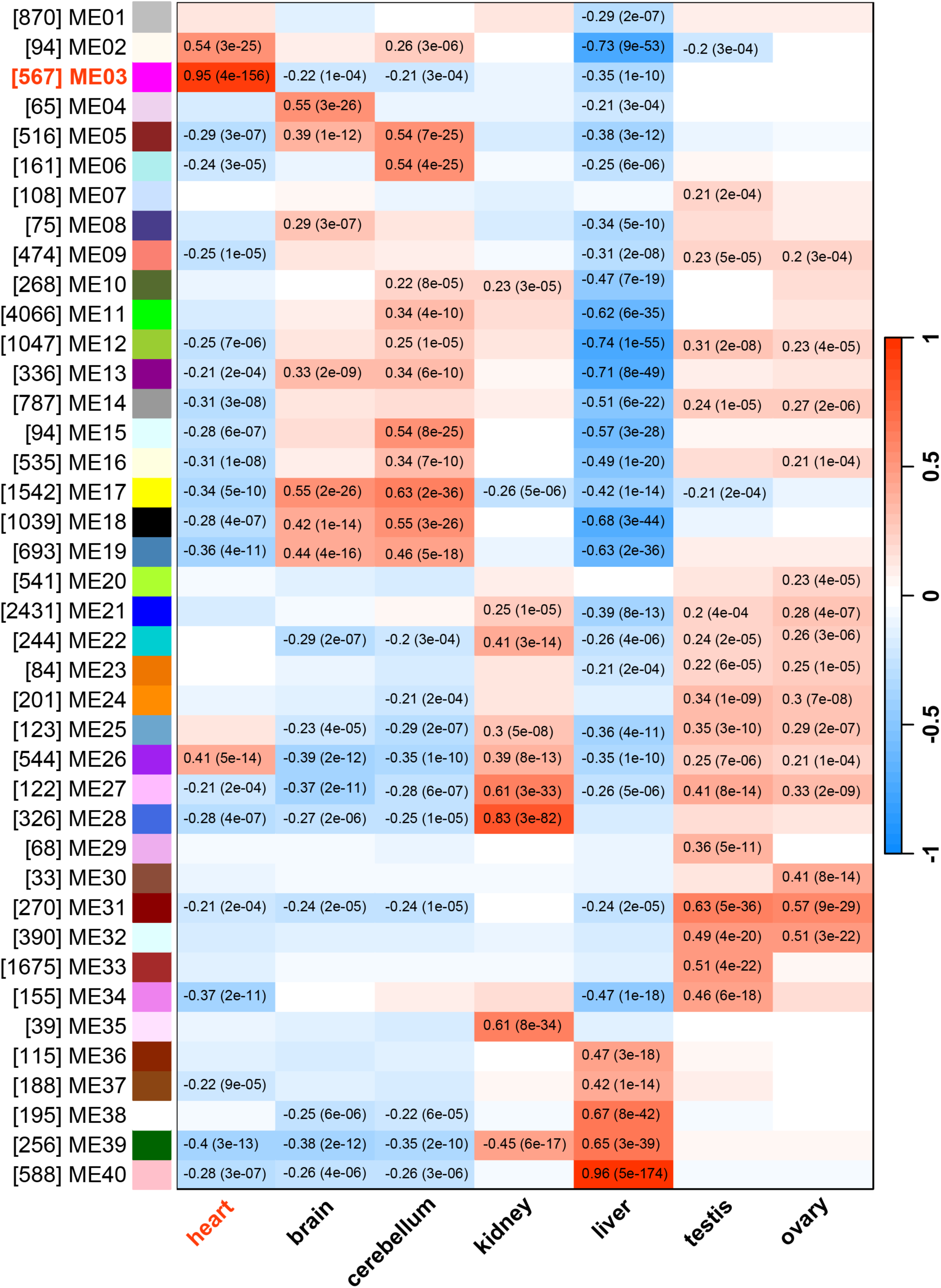
Module-Tissue Association Heatmap Across Seven Organ Systems. This heatmap helps identify gene modules critical for the development and function of specific tissues. It illustrates the correlation between the modules identified by WGCNA and the different tissue types across the developmental time series. Each row represents a module eigengene, color-coded according to the original module colors prefixed by “ME,” while each column corresponds to a specific tissue. Positive correlations are depicted in red, negative correlations in blue, and white indicates no correlation. The color intensity reflects the strength of the correlation, with numerical values in each cell displaying the correlation coefficient, followed by the p-value in parentheses for statistical significance. Only significant correlations (p < 0.001) are shown. Modules with high positive or negative correlations (> |0.6|) are considered tissue-specific, highlighting potential tissue-specific gene expression patterns. Module ME03 (magenta) notably shows the highest positive correlation (R = 0.95) with heart tissue, suggesting a strong link to heart development and function.

Hierarchical clustering of the modules demonstrated that the magenta module clustered most closely to the heart samples (**Supplementary Figure 8**). The magenta module (ME03), comprising 567 genes, demonstrated significant heart tissue specificity, with a positive correlation value of 0.95 and a p-value of 4 x 10^-156^. This module showed no significant positive correlations in the other organs (**Supplemental Data Table S3**). Focusing on the heart samples, we compared each module’s eigengene and developmental stage, using the age of the samples in post-conception (**Supplemental Data Table S4**). Modules with high positive correlations (in red) indicate increasing expression with developmental stage, while negative correlations (in blue) suggest decreasing expression over time (**Supplementary Figure 13**). The magenta module had a significant negative correlation (r = -0.50, p = 0.0002), clustering most closely with the mid-development (11–20 PCW) stage (**Supplementary Figure 14** and **Supplementary Figure 15**). In the magenta module, eigengene expression increased from conception, peaking around fetal mid-development (10-20 PCW) (**Figure 4**). Following this peak, expression levels gradually declined, reaching their lowest levels in the mature heart. CNVs impacting one or more of the 567 genes (425 protein-coding and 142 lncRNA genes) in the magenta module are found across all eight CHD phenotypes (**Supplemental Data Table S5**). In total, 154 (21%) of non-syndromic CHD patients had a CNV impacting the module (**Supplemental Data Table S6**).

**Figure 4.**
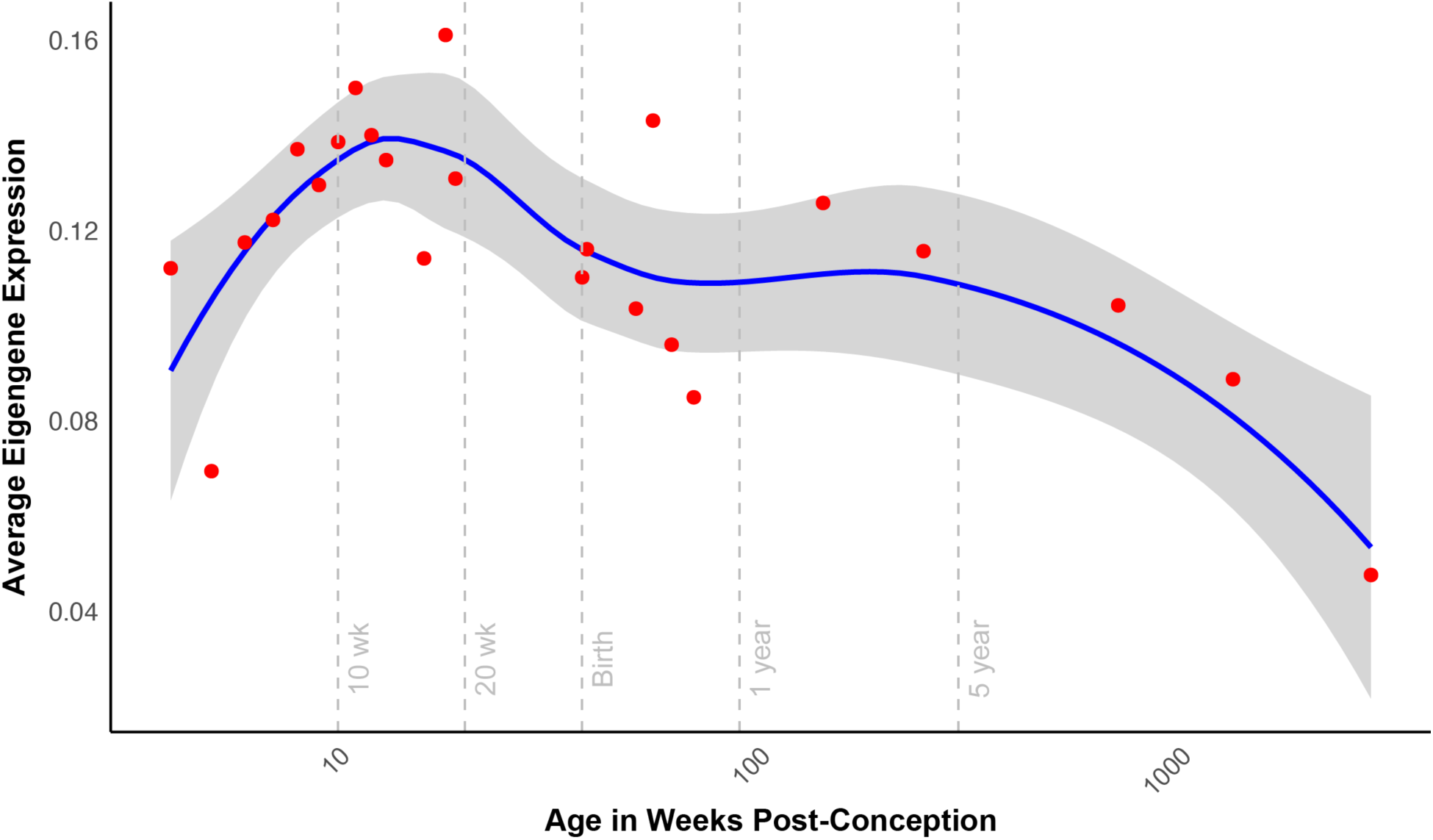
Magenta Module Eigengene Expression Across Developmental Stages in the Heart. The eigengene expression for the magenta module (ME03) is shown across various developmental stages in the heart, plotted by age (PCW). The red data points represent the average eigengene expression at each stage, with a LOESS-smoothed line (blue) fitted to highlight the trend over time. The gray shading indicates the 95% confidence interval, showing the range within which the trend line is expected to fall. Vertical dashed gray lines indicate key developmental milestones: 10 weeks, 20 weeks, birth (38 weeks), 1 year, and 5 years post-conception. These milestones mark significant transitions in heart development, providing context for the observed changes in eigengene expression. The expression level increases during early development, peaking between 10–20 PCW and gradually decreasing.

The functional associations of the magenta module were identified using gene ontology terms to reinforce its connection to the heart. Functional enrichment of GO ontology terms was performed using the 425 protein-coding genes within the magenta module. The analysis focused solely on protein-coding genes because lncRNA genes generally lack well-defined functional annotations in existing gene ontology databases, making them less suitable for GO enrichment analysis. After combining semantically similar terms, 159 GO terms were functionally enriched (**Supplemental Data Table S7**), with the top ten for each category illustrated in **Supplementary Figure 16**. These significant GO terms predominantly relate to muscle structure and function, highlighting critical processes such as muscle cell development, differentiation, and assembly. The identified cellular components include key structural elements like the sarcomere, myofibril, and contractile fibers. Additionally, molecular functions such as actin-binding, structural constituents of muscle, and electron transfer activity further emphasize the role of these genes in muscle contraction and energy transduction. Of the 159 enriched GO terms, 26 were related to the heart and the cardiovascular organ system (**Figure 5**). These terms included biological processes related to cardiac tissue muscle development (p = 4.41 x 10^-29^), heart contraction (p = 1.44 x 10^-21^), heart morphogenesis (p = 2.20 x10^-16^), cardiac chamber development (p = 2.79 x 10^-13^), vasculogenesis (p = 6.44 x 10^-05^) and embryonic heart tube development (p = 0.0037).

**Figure 5.**
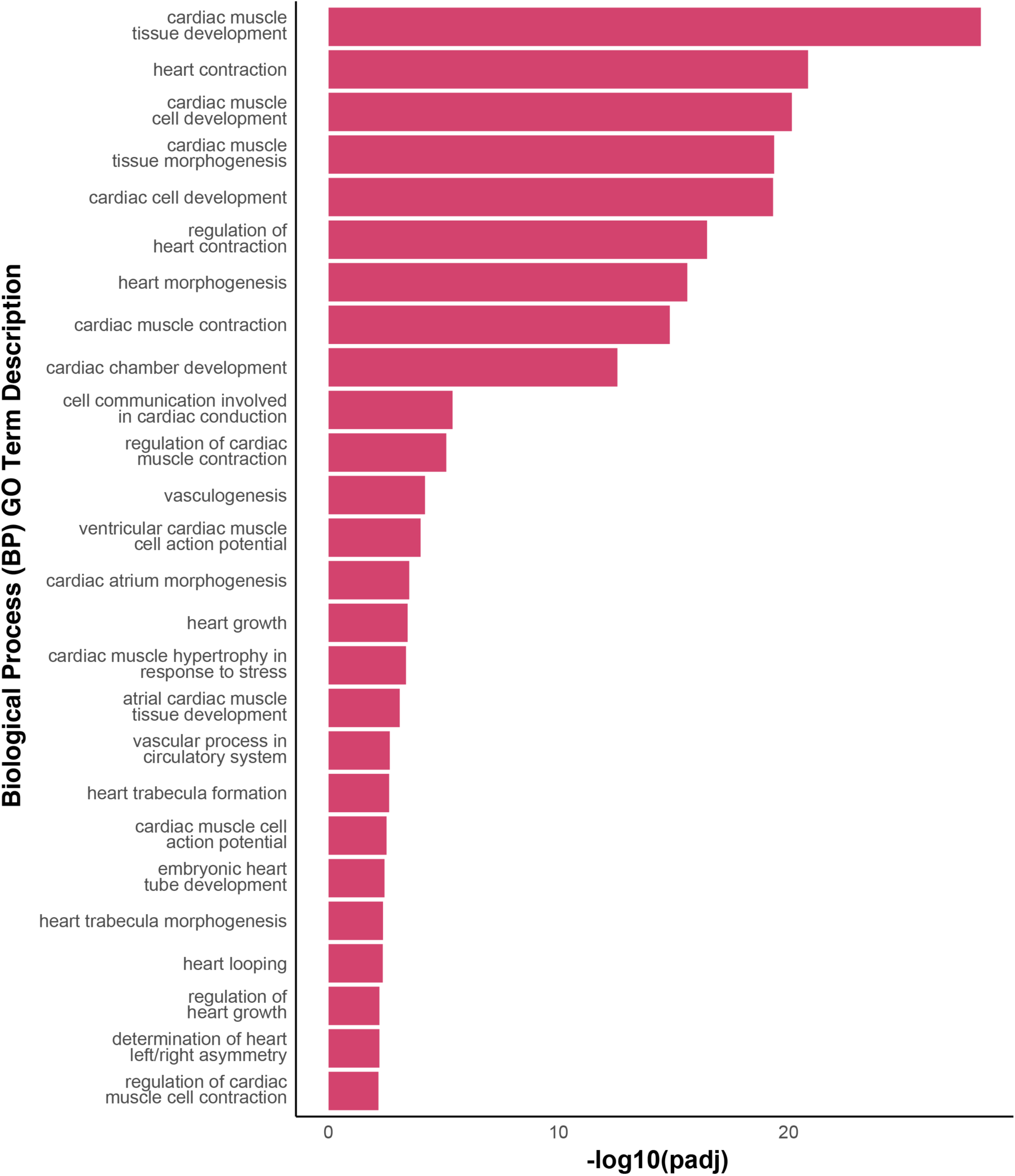
Functional Enrichment of Heart-Related Biological Processes in the Magenta Module. This bar chart presents 26 significantly enriched GO biological process (BP) terms related to the heart and cardiovascular system, derived from the protein-coding genes in the magenta module. Semantically similar terms were collapsed, and the most significant terms were selected based on their lowest adjusted p-values (Benjamini-Hochberg method), all below 0.05. Each bar represents the -log10 transformation of the adjusted p-values, indicating the relative statistical significance. The analysis highlights the magenta module’s enrichment for genes with key roles in heart development and function.

### Identification and Analysis of Hub Genes: Central Regulators of Heart-Specific Gene Networks

Genes with high module membership (**MM**), indicating strong connectivity within the module, and high gene significance (**GS**), indicating a strong association with the heart trait, were identified as hub genes (**Figure 6**). These genes are likely key drivers of the biological processes relevant to heart function and development. The correlation coefficients for the protein-coding genes (turquoise, r=0.968) and lncRNA genes (purple, r=0.972) both indicate strong positive correlation with p-value < 0.001. The correlation cutoff for module membership of r ≥ 0.80 led to the identification of 146 hub genes (25.7% of the total 567 genes in the module) (**Supplemental Data Table S8**). Within the hub genes, 18 were lncRNAs (12.3%, p=0.9326). Strikingly, eight of the nine known CHD genes in the module were within the hub, representing a highly significant enrichment (p=4.787 x 10^-06^). The lncRNA and known CHD hub genes and their corresponding MM and GS values are shown in **Table 1**. Our analysis identified 18 lncRNAs as potential clinical targets from the CCVM cohort of clinically validated CNVs associated with non-syndromic CHD (**Table 2**).

**Figure 6.**
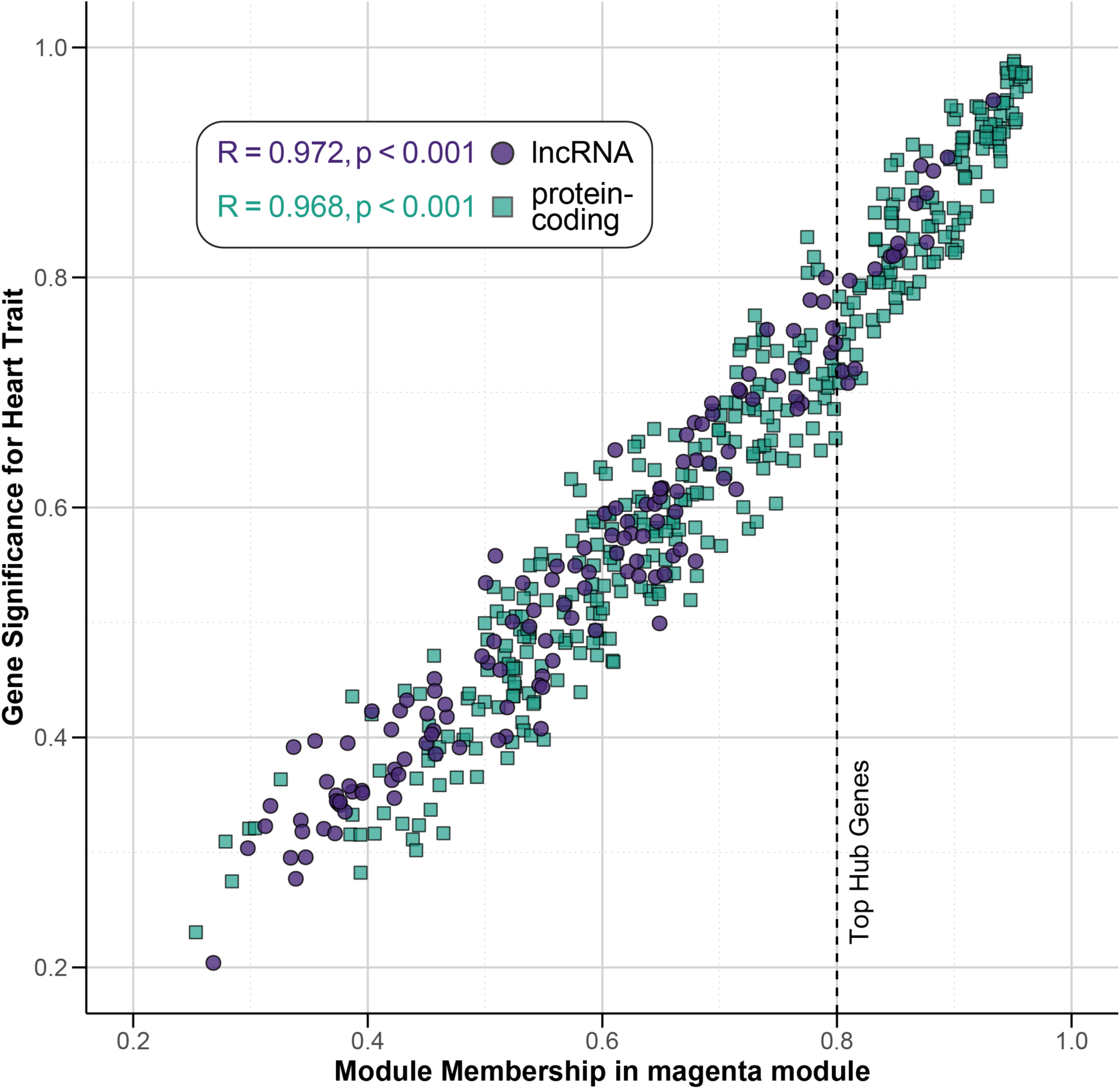
Identification of Hub Genes in the Magenta Module. This scatter plot illustrates the relationship between Module Membership (MM) and Gene Significance (GS) for genes within the magenta module (ME03). Module Membership reflects how strongly a gene is associated with the module eigengene, while Gene Significance measures the correlation between a gene’s expression and the heart trait. Genes are categorized into lncRNAs (purple circles) and protein-coding genes (teal squares). The top hub genes, which are highly connected within the module and strongly associated with the heart trait, are emphasized by a dashed vertical line at MM = 0.8. The plot also includes correlation (R) and p-values for both lncRNAs and protein-coding genes, calculated separately and displayed on the graph to indicate the strength and significance of the correlation between Module Membership and Gene Significance. This visualization is crucial for identifying lncRNA hub genes within the module, which may play essential roles in the heart-related biological processes associated with the module.

**Table 1.**
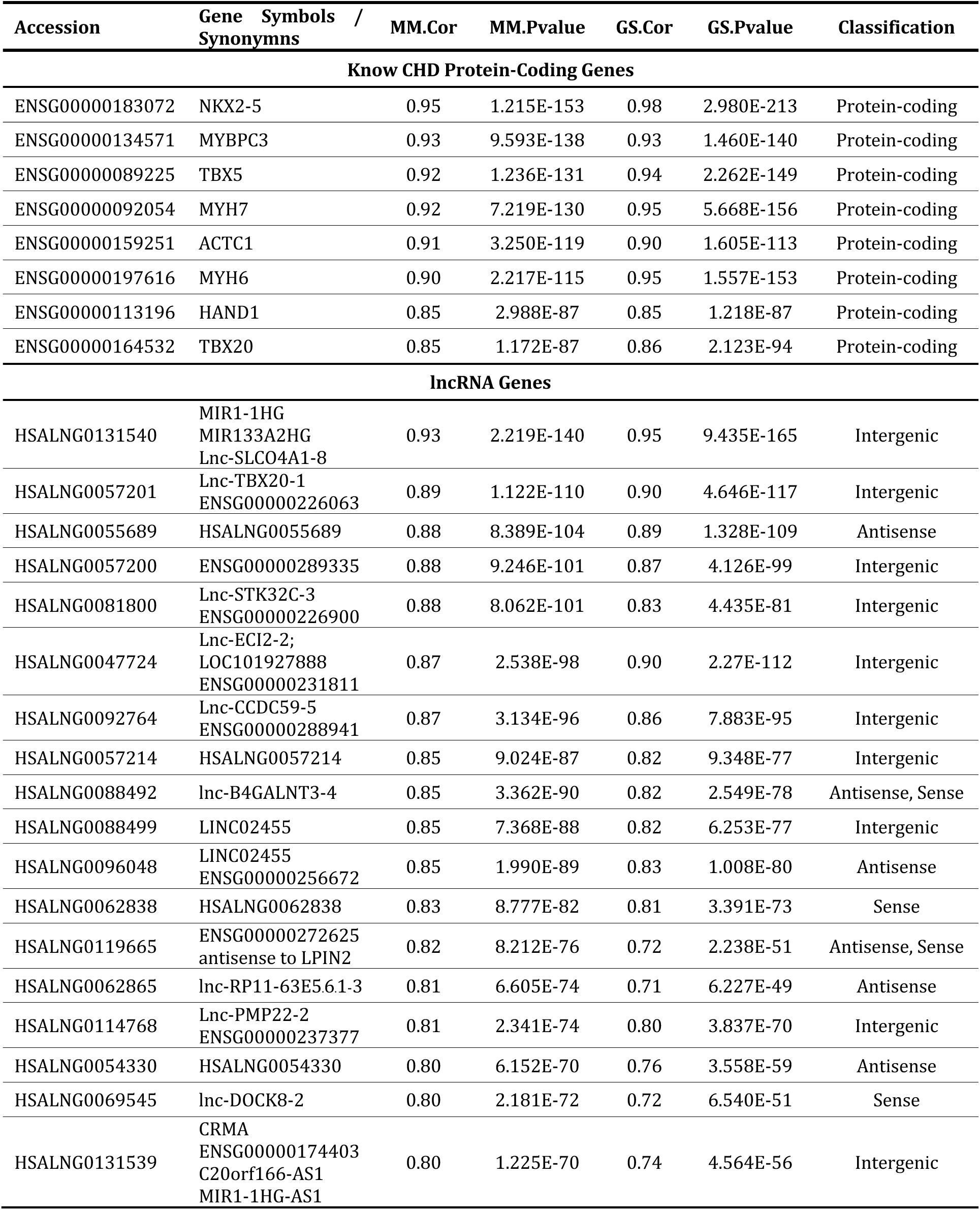
Known CHD and lncRNA Hub Genes from the Magenta Module (ME03). For each gene, the table provides the module membership correlation (MM.Cor) and its associated p-value, which indicates the strength of each gene’s association with the magenta module. Additionally, the table shows the gene significance correlation (GS.Cor) and its p-value, reflecting the correlation between each gene and heart-related traits. This summary highlights hub genes highly correlated with the module, including both protein-coding CHD genes and lncRNAs.

**Table 2:** lncRNA Hub Genes and Associated CNVs and Cardiac Phenotypes. Each lncRNA from the magenta module (ME03) is listed alongside the corresponding chromosomal region of the overlapping CNV (e.g., gain or loss) and the associated cardiac phenotype. The table also includes patient identifiers and CHD classification from the CCVM cohort who exhibit the relevant CNV, directly linking the genetic variations and observed heart conditions.

| LncBook Identifier | Overlapping CNV | Cardiac Phenotype | CCVM Patient(s) |
| --- | --- | --- | --- |
| HSALNG0047724 | 6p25.1 loss | CTD | 003-0293 |
| HSALNG0054330 | 10q23.1 gain<br>6q25.1 gain | CTD | 000-0369 |
| HSALNG0055689 | 7p22.3p22.2 loss | CTD | 002-0019 |
| HSALNG0057200<br>HSALNG0057201<br>HSALNG0057214 | 7p14.2 loss* | CTD | 003-0191 |
| HSALNG0062838 | 7q36.3 loss | LVOTO | 001-0182 |
| HSALNG0062865 | 8p23.3 loss<br>8p23.3 gain | CTD<br>LVOTO | 004-0065<br>003-0116 |
| HSALNG0069545 | 9p24.3 gain (2)<br>9p24.3 loss | CTD<br>RVOTO<br>LVOTO | 008-0053<br>001-0078<br>003-0375 |
| HSALNG0081800 | 10q26.3 loss (2) | RVOTO (2) | 002-0008<br>009-0009 |
| HSALNG0088492<br>HSALNG0088499 | 12p13.33 gain<br>2p16.3 loss | SD+LVOTO | 007-0016 |
| HSALNG0092764 | 12q21.31 loss<br>16p13.11 gain | SD | 003-0440 |
| HSALNG0096048 | 13q12.3 gain | CTD | 007-0002 |
| HSALNG0114768 | 2p21 gain<br>5p15.2 loss<br>17p12 gain | AVSD | 002-0007 |
| HSALNG0119665 | 18p11.32p11.31 loss<br>18p11.32p11.31 gain | APVR<br>SD | 005-0036<br>000-0224 |
| HSALNG0131539<br>HSALNG0131540 | 20q13.33 gain | SD+LVOTO | 001-0285 |

\*This CNV also contains the known CHD gene TBX20

The relationships within the magenta module were further explored through co-expression network visualization (**Figure 7**). To enhance the clarity of the network, we applied a stringent filter to the edges, retaining only those with a TOM value greater than 0.30. This filtering resulted in a network comprising 67 nodes with high module membership (MM.Cor ≥ 0.80) and strong co-expression interaction. Known CHD genes in the module were included in the network regardless of their edge threshold interaction value. As a result, the network included 8 CHD genes, 7 lncRNA genes, and 52 protein-coding genes. Notably, all genes in this network displayed a co-expression interaction with *NKX2-5*, a critical regulator of heart development.

**Figure 7.**
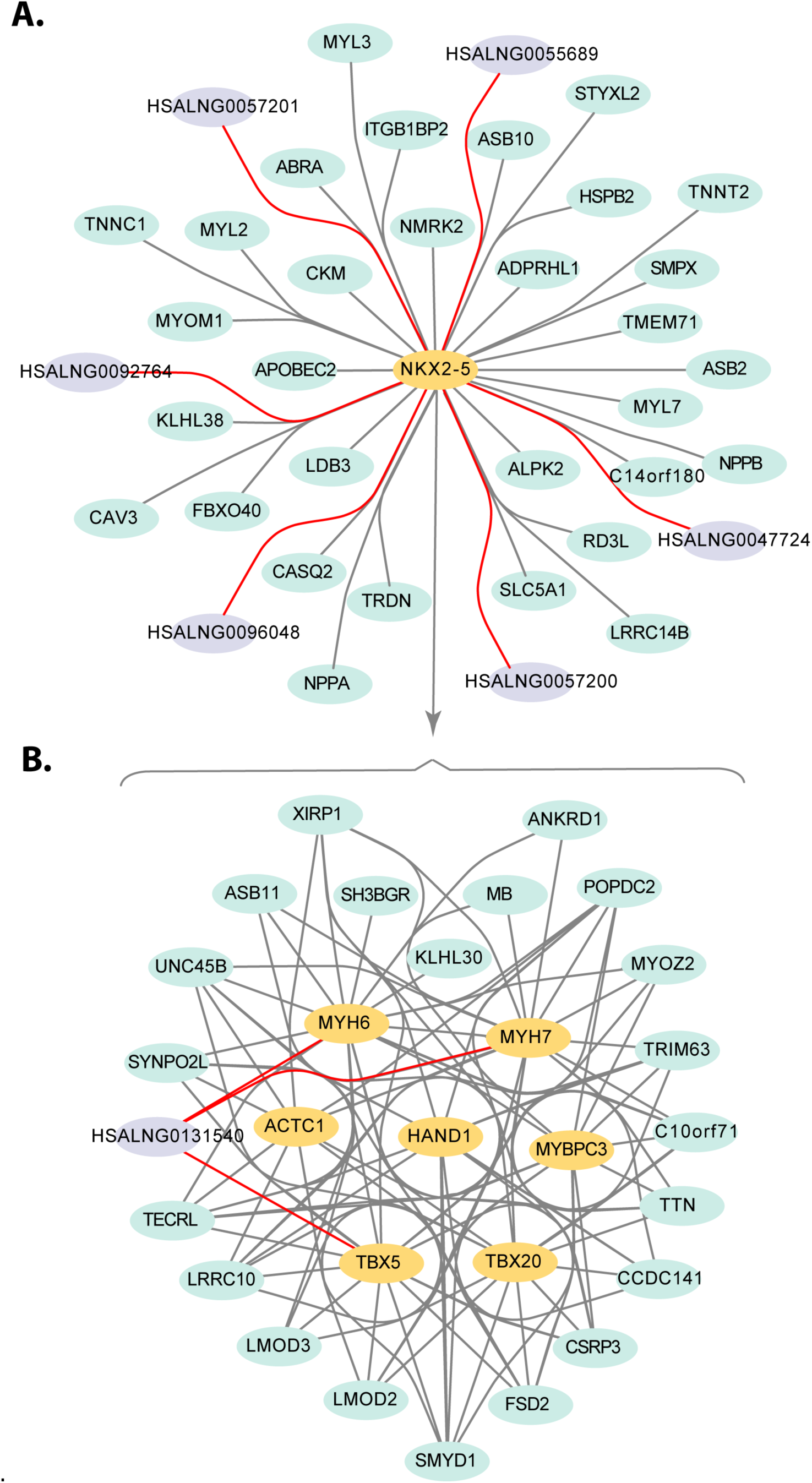
Gene Network Visualization of Hub Genes in the Magenta Module. This figure visualizes the co-expression network of hub genes within the magenta module (ME03), generated using Cytoscape. Nodes represent hub genes, color-coded by gene type: gold for known CHD genes, purple for lncRNAs, and light green for protein-coding genes. Edges indicate co-expression interactions between genes based on the TOM matrix, with a threshold of TOM > 0.30. Hub genes are defined as those with high module membership (R ≥ 0.80). Co-expression edges connecting to lncRNA genes are highlighted in red. The network is divided into two panels for clarity: (**A**) Seven of the eighteen CNV-lncRNA hub genes are co-expressed with *NKX2-5,* a critical transcription factor for early heart development. (**B**) A highly interconnected network of known CHD genes and other protein-coding genes, all co-expressed with *NKX2-5*. For clarity, redundant edges were removed and indicated with an arrow and curly brace.

The Gini index of over 36,000 genes from 31 tissues, developed using independent human heart RNA-seq and GTEx datasets, was mapped to the hub genes.^28^ The Gini coefficient with respect to the distribution of expression levels across samples identifies genes likely important in heart development by highlighting those with expression patterns specific to the heart during embryogenesis. VanOudenhove *et al.* proposed that candidate CHD genes have a Gini coefficient greater than 0.5 and, of the tissues studied, the gene must be maximally expressed in the embryonic heart. Within the hub genes identified in the magenta module, all eight known CHD genes and seven of the 18 lncRNAs genes had Gini scores from their study (**Table 3**). Four of the seven lncRNA candidates met the CHD gene candidate criteria: *lnc-TBX20-1* (Gini = 0.959), *lnc-STK32C-3* (Gini = 0.867), *CRMA* (Gini = 0.662), and *lnc-DOCK8-2* (Gini = 0.631). As was observed in the known CHD genes MYH6 and MYH7, the other candidate lncRNA genes had high expression in adult heart and muscle, or had a Gini score below 0.5, thus not meeting VanOudenhove *et al.’s* suggested criteria for a CHD candidate gene.

**Table 3:** Gini Coefficient Analysis of Hub Genes. This table presents the Gini coefficients for hub genes identified in the magenta module (ME03). The table includes hub genes that are either known CHD genes or lncRNAs. The Gini coefficient, a measure of tissue-specific gene expression, is calculated for each gene, with higher values indicating tissue-specific expression. Genes with a Gini coefficient > 0.5 and maximal expression in the embryonic heart are considered to be candidate CHD genes. Out of 8 known CHD hub genes, five were found to have significant embryonic heart-specific expression (Gini coefficient > 0.5). Among the 18 lncRNA hub genes, 7 of the 9 with Ensembl accessions have a Gini coefficient, and five are maximally expressed in the embryonic heart. Of these, 4 lncRNAs exhibit significant heart-specific expression with a Gini coefficient greater than 0.5.

| Accession | Symbol | Type | Gini Coefficient | Tissue with max expression | Max Expression |
| --- | --- | --- | --- | --- | --- |
| ENSG00000226063 | Lnc-TBX20-1<br>(HSALNG0057201) | lncRNA | 0.959 | Embryonic Heart | 167 |
| ENSG00000164532 | TBX20 | CHD | 0.948 | Embryonic Heart | 5117 |
| ENSG00000197616 | MYH6 | CHD | 0.936 | Heart | 323262 |
| ENSG00000183072 | NKX2-5 | CHD | 0.935 | Embryonic Heart | 9467 |
| ENSG00000134571 | MYBPC3 | CHD | 0.934 | Heart | 91682 |
| ENSG00000231811 | Lnc-ECI2-2<br>(HSALNG0047724) | lncRNA | 0.93 | Heart | 85 |
| ENSG00000174407 | MIR1-1HG<br>(HSALNG0131540) | lncRNA | 0.918 | Muscle | 5447 |
| ENSG00000159251 | ACTC1 | CHD | 0.915 | Embryonic Heart | 290948 |
| ENSG00000092054 | MYH7 | CHD | 0.905 | Muscle | 457406 |
| ENSG00000113196 | HAND1 | CHD | 0.889 | Embryonic Heart | 2246 |
| ENSG00000089225 | TBX5 | CHD | 0.873 | Embryonic Heart | 5400 |
| ENSG00000226900 | Lnc-STK32C-3<br>(HSALNG0081800) | lncRNA | 0.867 | Embryonic Heart | 567 |
| ENSG00000174403 | CRMA / MIR1-1HG-AS1<br>(HSALNG0131539) | lncRNA | 0.662 | Embryonic Heart | 2167 |
| ENSG00000226403 | lnc-DOCK8-2<br>(HSALNG0069545) | lncRNA | 0.631 | Embryonic Heart | 398 |
| ENSG00000272625 | antisense to LPIN2<br>(HSALNG0119665) | lncRNA | 0.251 | Embryonic Heart | 81 |

## Discussion

We applied WGCNA to cluster CNV-lncRNAs using a similar approach to that described for the heart and kidney.^34,46^ CNV-associated lncRNAs were identified in clinically reported CNVs from a cohort of patients with isolated CHD. Utilizing human organ transcriptomic time-series data for the development of seven major organs, we clustered gene expression of both protein-coding and CNV-lncRNA genes into modules. These modules were then correlated with specific organs, allowing us to identify a module of co-expressed genes highly specific to the developing heart. We identified eighteen CHD-candidate lncRNA genes within this heart-specific module (the magenta module, ME03). Seven of these eighteen lncRNAs had significant co-expression relationships with known CHD genes, and four were confirmed as candidate CHD genes in an orthogonal data set examining gene expression patterns specific to the heart during embryogenesis. Our study offers a well-documented and reproducible programming pipeline that applies a widely used method in a novel way to identify candidate lncRNAs within a disease-specific context and predict their functions. This approach empowers researchers to systematically study CNV-lncRNAs, providing a comprehensive toolset for future investigations. Moreover, our work expands the analysis of CNV-lncRNAs by utilizing a larger and more recent dataset of clinically confirmed CNVs from CHD patients in the CCVM dataset, further enhancing the scope and applicability of the research.

Clinical genetics in CHD primarily emphasizes well-known protein-coding genes, as evidenced by resources such as the high-confidence CHDgene list.^7^ However, our analysis of the CCVM cohort revealed that only 72 of the 953 clinically reported CNVs (7.6%) affected one of these known CHD genes, suggesting that many potential pathogenic factors remain unidentified. By contrast, 656 of the 953 clinically reported CNVs (68.8%) contain one or more lncRNAs expressed in the heart, i.e., CNV-lncRNAs, underscoring the need to consider the non-coding genome as a crucial factor in the genetic etiology of CHD.

Through WGCNA, we identified a highly heart-correlated module (the magenta module, ME03, **Figure 3**) consisting of 567 genes with significant heart tissue specificity, clustering most closely with the mid-development stage (11–20 PCW). No significant positive correlations for this module were observed in other organs. CNVs affecting these genes were present across all eight CHD phenotypes, with 21% of non-syndromic CHD patients carrying a CNV impacting the module. In this module, eigengene expression increased steadily from conception, peaking during fetal mid-development (10–20 PCW) (**Figure 4**). After this peak, expression levels gradually declined, reaching their lowest point in the mature heart. This expression pattern suggests that the genes within the magenta module are most active during early to mid-heart development, a critical period for structural refinement, including chamber septation, conduction system development, and the initiation of blood circulation. Functional enrichment analysis of the protein-coding genes in this module identified multiple enriched GO terms directly linked to heart and cardiovascular development, highlighting processes such as cardiac muscle development, heart contraction, and morphogenesis (**Figure 5**).

Additionally, the analysis revealed significant enrichment of genes involved in mitochondrial function, energy production, and metabolic processes, underscoring the critical role of cellular energy dynamics in heart development (**Supplemental Data Table 7** and **Supplemental Figure 16**). Given the essential role of mitochondria in providing the energy required for cardiac muscle contraction and overall cardiovascular function, this enrichment of mitochondrial and metabolic genes suggests that dysregulation in energy metabolism may contribute to CHD pathogenesis. Overall, the developmental gene expression pattern and the identified gene ontology terms suggest that genes in this module likely play pivotal roles in heart development and may serve as candidate genes for CHD.

Within the module of genes highly correlated with the developing heart, we identified eighteen highly connected lncRNA hub genes strongly correlated with the module eigengene (**Figure 6**). These genes are likely key drivers of the biological processes relevant to heart function and development. However, the lack of Ensembl accession numbers for many of our candidate lncRNAs limited our search for external information. Notably, only nine of these lncRNAs possess recognized Ensembl identifiers; the other nine were putative lncRNAs identified by LncBook. This highlights significant constraints in current gene annotation methods, which are often too stringent for ncRNA. For instance, Ensembl’s criteria exclude intergenic lncRNAs with open reading frames that exceed 35% of their length or contain protein domains.^47^ Consequently, our analysis and subsequent discussions were confined to those lncRNA candidates with Ensembl identifiers, limiting the full exploration of the potential roles of the remaining putative lncRNAs in heart development and emphasizing the need for more inclusive gene annotation frameworks for non-coding RNAs.

Seven of the eighteen CNV-lncRNA hub genes were co-expressed with *NKX2-5*, a well-known transcription factor critical for early heart development: HSALNG0057201 (Lnc-TBX20-1), HSALNG0057200 (ENSG00000289335), HSALNG0055689, HSALNG0092764 (Lnc-CCDC59-5), HSALNG0096048 (ENSG00000256672), HSALNG0047724 (Lnc-ECI2-2), and HSALNG0131540 (MIR1-1HG) (**Figure 7A**). *NKX2-5* is expressed in the first and second heart fields to maintain chamber-specific identities.^48^ Pathogenic variants in *NKX2-5* are associated with a variety of CHDs, including atrial^49^ and ventricular SDs^50^, AVSDs^51^, and others^52,53^. Notably, HSALNG0131540 or *MIR1-1HG* was also significantly co-expressed with well-established CHD genes, *TBX5*^54,55^, *MYH6*^56,57^, and *MYH7*^58,59^ (**Figure 7B**). The co-expression of these lncRNAs with *NKX2-5* and other known CHD genes underscores their potential roles in heart development and cardiac function. This suggests that they may be involved in similar developmental pathways and could contribute to the pathogenesis of congenital heart defects.

Four of our CNV-lncRNA hub genes—*lnc-TBX20-1*, *lnc-STK32C-3*, *CRMA*, and *lnc-DOCK8-2*— were also highly ranked as candidate CHD genes in an external validation (**Table 3**). These genes met the criteria of having the highest expression in the embryonic heart across 32 different organs and tissues, along with a significant Gini index, indicating strong heart-specificity.^28^ Established CHD genes that meet the Gini index criteria include *TBX20* (Gini=0.948) and *NKX2-5* (Gini=0.935). Unexpectedly, the three known CHD genes—*MYBPC3*, *MYH6*, and *MYH7*—were excluded from VanOudenhove’s candidate gene set because their expression extends beyond the embryonic heart into the adult heart and muscle. Despite the Gini index limitations to score all known CHD genes, this index did confirm *lnc-TBX20-1*, *lnc-STK32-C*, *CRMA*, and *lnc-DOCK8-2*—as potential contributors to heart-specific gene regulation and strengthens their candidacy for further investigation in CHD research.

The intergenic lncRNA *lnc-TBX20-1* (HSALNG0057201) was particularly interesting due to its high Gini index and proximity to the known CHD gene *TBX20*^60^. Of the known CHD genes and CNV-lncRNAs, *lnc-TBX20-1* had the highest Gini index (0.959) and peak expression in the embryonic heart, making it a strong CHD candidate lncRNA. The CNV associated with *lnc-TBX20-1* is a 350 Kb loss of 7p14.2 in a patient with CTD. This region contains the known CHD gene, *TBX20*, and two other lncRNA candidates, HSALNG0057200 and HSALNG0057214. The location of lncRNAs is important as lncRNAs are known to regulate nearby genes.^10^ While it is not possible to determine whether the effects of this pathogenic CNV are due to the loss of *TBX20* or the lncRNA genes, the co-expression of *lnc-TBX20-1* and HSALNG0057200 with *NKX2-5* is particularly intriguing. *NKX2-5* and *TBX20* are transcription factors that interact in heart development regulation.^61^ This raises the possibility that these lncRNAs may have regulatory roles in connection with *TBX20* and potentially *NKX2-5*, though experimental evidence is still needed to support this hypothesis.

lncRNAs function in processes ranging from chromatin modification to nuclear organization and are emerging as critical players in cardiovascular development and disease. However, their complexity often extends beyond the traditional definitions of non-coding genes, as exemplified by *lnc-STK32C-3*’s (HSALNG0081800) coding potential.^13^ The CNVs associated with *lnc-STK32C-3* are two separate 2.9 MB deletions of 10q26.3 observed in two CCVM patients with RVOTO. We identified this CNV-lncRNA as a candidate CHD gene with the highest module membership in the magenta module, indicating specificity in cardiac development. This gene was also indicated as a candidate CHD gene by the Gini index with highly significant Gini index of 0.867, and is most highly expressed in the embryonic heart across the 32 organs and tissues transcriptionally profiled.^28^ We are the first to suggest that this lncRNA is involved in the initial formation of the heart. Previous research suggests that *lnc-STK32C-3* is abundantly expressed in ischemic hearts and it is suggested to be involved in myocardial injury.^14^ This aligns with the broader understanding that genes can turn on and off in response to developmental cues and cardiac injury, as shown by the protein-coding gene *VEGF-B*.^62^ Therefore, if protein-coding genes can have dual roles under different conditions expressed by the heart, it is plausible that lncRNAs can, too. Lastly, while it is known that *lnc-STK32C-3* encodes microproteins and secretes them in the adult heart,^27^ whether it can also perform this function during cardiac development remains to be explored.

*CRMA* is a known regulator of cardiac differentiation and has been experimentally validated as important for heart development.^24,63^ Variants affecting *CRMA* are therefore likely pathogenic and could help explain the CHD phenotype. The CNV associated with *CRMA* is a 2.9 MB gain in the 20q13.3 region in a patient with SD and LVOTO. In addition to *CRMA*, this CNV impacts the known CHD gene *GATA5*^64,65^ and the lncRNA *MIR1-1HG*^24^. *MIR1-1HG* is the precursor of two cardiogenic miRNAs, known as *MIR1-1* and *MIR-133a2*. *GATA5* has only been implicated in CHD as a loss-of-function mutation in humans,^64^ whereas in our cohort patients with 20q13.3 is a gain-of-function mutation. When CRMA is expressed, it downregulates *MIR1-1* and *MIR133a2,* pushing embryonic stem cells into a mature cardiac fate.^24,63^ While the knockdown of *CRMA* promotes cardiac differentiation through increased expression of MIR-133a2, MIR-133a1 targets RBPJ to inhibit the NOTCH pathway.^24^ We hypothesize that in a gain-of-function mutation CRMA may suppress the cardiac miRNAs expression beyond normal levels, leading to dysregulation of the NOTCH pathway causing cardiac malformations. Together, these networks of genes are required for the coordinated regulation of cardiogenic differentiation in embryonic stem cells.

Three CNVs impacting 9p24.3 were associated with *lnc-DOCK8-2:* a 305 Kb duplication in a patient with RVOTO, a 354 Kb duplication in a patient with CTD, and a 100 Kb deletion in a patient with LVOTO. While the CNVs do not impact any known CHD genes, in addition to the lncRNA, these smaller CNVs impact two protein-coding genes, *DOCK8* and *KANK1*. *DOCK8* is a known Mendelian disease gene, resulting in a rare autosomal recessive immunologic disorder characterized by recurrent staphylococcal infections of the skin and respiratory tract.^66^ *KANK1* is an essential regulator of the actin cytoskeleton, similar to Rho GTPases, which also regulate actin dynamics through Rho-kinase (ROCK), and is required for myogenic differentiation.^67^ Rho GTPase signaling is involved in cardiac looping and early development of the heart, and dysregulation of this pathway may lead to structural heart defects.^68^ Whether *lnc-DOCK8-2* regulates either of these genes is unknown, and like the majority of CNV-lncRNAs that we identified, further studies are required.

Reproducible bioinformatics methods and well-documented software are still significant barriers to advancing biomedical research, often hindering the ability to replicate and build upon existing studies. To address this challenge, we developed comprehensive and well-documented R notebooks that streamline our analytical workflow and fully automate the generation of all tables and figures, enabling easy replication and modification of the entire pipeline. This approach provides a robust and adaptable framework standardizing co-expression analysis to CNV-lncRNAs that can be applied to other genomic datasets, facilitating broader investigations into the roles of lncRNAs in tissue-specific and disease-specific contexts. As annotations of non-coding RNAs continue to improve and new genomic studies of CHD patients emerge, our methods can be reused and extended to uncover additional candidate genes and pathways involved in CHD.

### Conclusion

In this study, we identified and characterized previously unrecognized lncRNAs associated with well-established protein-coding CHD genes, confirming prior findings while revealing new potential links, such as the involvement of lnc-DOCK8-2 and HSALNG0096048 in our CHD cohort. By applying the advanced bioinformatics approach of WGCNA, we uncovered insights into the complexities of CNV-associated lncRNAs within a CHD context. These findings emphasize the potential pathogenic roles of non-coding regions, which are often overlooked by traditional genetic studies. Our work highlights the significant, yet underexplored, contribution of lncRNAs to the genomics of CHD, opening new pathways for diagnosing and understanding the genetic basis of CHD.

In addition to these discoveries, we have made our fully documented R code publicly accessible, enabling other researchers to replicate and build upon our findings. This transparency facilitates further investigation into the uncharted territory of lncRNAs, their critical roles in heart development, and their probable link to CHD. As we continue to advance our understanding of non-coding RNAs and their role in CHD, we anticipate that this research will ultimately lead to better diagnostic tools and improved outcomes for individuals affected by CHD. As genome sequencing is incorporated into the standard of care for patients with critical CHDs, we encourage clinical labs to consider reporting CNVs and single nucleotide variants impacting lncRNAs.

## Acknowledgments

We thank the Nationwide Children’s Hospital Foundation and The Abigail Wexner Research Institute at Nationwide Children’s Hospital for generously supporting this work. These funding bodies had no role in the study’s design, collection, analysis, and interpretation of data, nor in writing the manuscript. Thank you to Thomas Hohmann, who contributed to data analysis early on in the project, and Dhruv Prasad, who tested the reproducibility of the CNV-lncRNA pipeline.

## Ethics Approval and Consent to Participate

This study exclusively employed publicly available deidentified datasets and thus did not involve direct human subjects research. For the CCVM data, collection of the clinical CMA reports utilized a waiver of informed consent approved by each clinical center’s Institutional Review Board. In conducting this analysis, we strictly adhered to the datasets’ terms of use, access, and distribution as outlined by their respective sources. We ensured that our research methods and objectives were aligned with the ethical guidelines for research and data use, including respecting privacy, intellectual property rights, and the integrity of the data.

## Data Availability

The CCVM dataset, used for identifying CNV-lncRNA interactions, is accessible through the CCVM Consortium. The RNA-seq data, including expression levels of lncRNAs, is available from LncExpDB at the lncRNA Expression Database (https://ngdc.cncb.ac.cn/lncexpdb/). The sample metadata was downloaded from Array Express (https://www.ebi.ac.uk/biostudies/arrayexpress/studies/E-MTAB-6814). The LncBook GTF file, used to map CNV-lncRNAs to the GRCh38 genome build and quantify gene expression, is available at LncBook (http://bigd.big.ac.cn/lncbook).

The CHDgene database, which provided a curated list of genes associated with CHD, is available at https://chdgene.victorchang.edu.au. The Gini Index data used for tissue-specificity analysis, published by the Cotney Lab, is available at https://cotneylab.cam.uchc.edu.

The source code used for CNV-lncRNA mapping, preprocessing, and network construction is available under an open-source license on GitHub (https://github.com/nch-igm/lncCHDNet/). This ensures that other researchers can freely access, modify, and implement the code in their respective analyses. The code is also stably archived on Zenodo at https://doi.org/10.5281/zenodo.13799847 to ensure long-term availability and reproducibility.

