## Supplementary Methods for "Identification of Long Non-coding RNA Candidate Disease Genes Associated with Clinically Reported CNVs in Congenital Heart Disease"

[†] Cytogenomics of Cardiovascular Malformations (CCVM) Consortium – see **Supplemental Methods** for the complete list of Contributing Authors.

**Short title**: CNVs impacting lncRNAs in congenital heart disease

***Corresponding Author:** Peter White, Ph.D., Battelle Endowed Chair for Quantitative and Computational Medicine, The Office of Data Sciences, The Abigail Wexner Research Institute at Nationwide Children's Hospital, 575 Children's Crossroad, Columbus, OH 43215. USA.

**Cytogenomics of Cardiovascular Malformations (CCVM) Consortium Contributors**

<table>
<tr><th>Name</th><th>Contributing Organization</th></tr>
<tr><td>Stephanie M. Ware</td><td rowspan="5">Department of Pediatrics, Indiana University School of Medicine, Indianapolis, Indiana, United States of America<br>Department of Medical and Molecular Genetics, Indiana University School of Medicine, Indianapolis, Indiana, United States of America</td></tr>
<tr><td>Benjamin J. Landis</td></tr>
<tr><td>Lindsey R. Helvaty</td></tr>
<tr><td>Gabrielle C. Geddes</td></tr>
<tr><td>Jennelle C. Hodge</td></tr>
<tr><td>Kim L. McBride</td><td rowspan="2">Center for Cardiovascular Research, Nationwide Children's Hospital, Columbus, Ohio, United States of America.<br>Department of Pediatrics, The Ohio State University College of Medicine, Columbus, Ohio, United States of America</td></tr>
<tr><td>Vidu Garg</td></tr>
<tr><td>Cecilia W. Lo</td><td rowspan="2">Department of Developmental Biology, University of Pittsburgh, Pittsburgh, Pennsylvania, United States of America</td></tr>
<tr><td>Svetlana A. Yatsenko</td></tr>
<tr><td>Jiuann-Huey Ivy Lin</td><td>Department of Critical Care Medicine, University of Pittsburgh, Pittsburgh, Pennsylvania, United States of America</td></tr>
<tr><td>Stephanie Burns Wechsler</td><td>Department of Pediatrics, Emory University School of Medicine, Atlanta, Georgia, United States of America</td></tr>
<tr><td>Seema R. Lalani</td><td>Department of Molecular and Human Genetics, Baylor College of Medicine, Houston, Texas, United States of America</td></tr>
</table>

**LncExpDB**

LncExpDB reprocessed the original organ developmental time series RNA-seq dataset by starting with FASTQ sequence reads, aligned against the GRCh38/hg38 human genome assembly using the STAR aligner (v2.7.1a). Transcript abundances were quantified with Kallisto (v0.46.1), and transcript per million (**TPM**) values were calculated for each transcript. The analysis pipeline integrated transcript annotations from LncBook (v1.2) and GENCODE (v33), generating a high-confidence list of lncRNAs. For samples with biological replicates, TPM values were averaged. Gene-level aggregation was performed using GffCompare, linking transcripts with shared exonic sequences on the same strand to form gene-level expression profiles. This approach resulted in a comprehensive expression profile that includes 101,293 lncRNAs and 19,957 protein-coding genes. The LncExpDB processing pipeline generated abundance matrices (read counts, CPM, FPKM, and TPM) normalized using the TMM method. LncRNAs with a maximum TPM value of less than 1.0 in any biological condition were classified as unexpressed.

**WGCNA**

***Step 1: Calculate Topological Overlap Matrix (TOM)***

The Topological Overlap Matrix (TOM) measures the similarity between genes based not only on their direct connections but also on their shared connections with other genes, thereby capturing the network's topological structure. TOM is computed from the adjacency matrix, representing the pairwise connection strengths between genes. The adjacency matrix is derived from the expression data using a soft-thresholding power: entry $a_{i,j}$ is the absolute value of the Pearson correlation between expression profiles of genes $i$ and $j$, raised to the power of β.

We evaluated a range of powers to ensure an optimal balance between achieving a scale-free topology and maintaining network connectivity (**Supplemental Figure 5**). While a power of 16 is the first to cross the strict $R^2$=0.80 threshold, powers in the range of 6-15 are very close to this value. Often, a slightly lower $R^2$ (e.g., around 0.75-0.79) can still be considered adequate, particularly if it results in a more interpretable network. As such, a power value of 9 ($R^2$=0.76) was selected to offer a good balance, providing sufficient connectivity for meaningful module detection while still maintaining an approximate scale-free topology. This approach allows for a more detailed examination of gene interactions and the potential regulatory mechanisms involved in disease pathology. TOM then refines the adjacency matrix by considering the overlap in connections.

The TOM between genes $i$ and $j$ is calculated using the following equation:

$$TOM_{i,j} = \frac{a_{i,j} + \sum_{u \neq i,j} a_{i,u} . a_{u,j}}{min(k_i, k_j) + 1 - a_{i,j}}$$

Where $k_i$ and $k_j$ denote the connectivity of genes $i$ and $j$, respectively:

$$k_i = \sum_{u \neq i} a_{i,u}$$

The term $\sum_{u \neq i,j} a_{i,u} . a_{u,j}$ measures the number of shared neighbors between $i$ and $j$ in the adjacency graph. The connectivity terms, $k_i$ and $k_j$, represent the total strength of neighbor connections for gene $i$ in the adjacency graph. The term $min(k_i, k_j)$ normalizes the TOM value by considering the connectivity of the less connected gene. That connectivity term contains an addition of $a_{i,j}$, and finally, $1 - a_{i,j}$ removes that term and adds 1 to regulate the denominator. By considering both direct and shared connections, the TOM equation helps identify clusters of highly interconnected genes, which can be used to infer functional relationships and identify gene modules in co-expression networks.

The TOM was generated using the TOMSimilarityFromExpr() function from the WGCNA package to the 21,925 genes (3,608 lncRNA and 18,317 protein coding) with the parameters *power=9* and *networkType=''*.

***Step 2: Detect Network Modules (Clusters)***

In WGCNA, genes are grouped into co-expression modules based on their correlation patterns. Each module is assigned a unique identifier in the form of a color, which provides a visual and intuitive way to distinguish between different modules. These colors are arbitrary and serve as labels, but they remain consistent throughout the analysis, making it easy to track modules across different plots and analyses. For example, genes assigned to the “magenta” module exhibit similar expression patterns, while those in the “green” module form a distinct co-expression group. This color-based labeling system facilitates the interpretation of network topology and gene clustering results.

Hierarchical clustering is performed on the distance matrix derived from the TOM, specifically using $1 - TOM$ to measure gene-gene dissimilarity (**Supplemental Figure 6**). Modules, or groups of co-expressed genes, are identified using the dynamic tree cut method, which segments the hierarchical tree into distinct branches representing different modules. The dynamic tree cut method (*cutreeDynamic* function in WGCNA) is applied to this dendrogram for module identification, utilizing parameters to optimize the detection of biologically relevant modules. The "*hybrid*" method, which is the default in WGCNA, offers greater flexibility and robustness, especially in complex datasets where a strict hierarchical approach might miss important relationships. Additional options included *minClusterSize=30* to ensure that each module contains at least 30 genes, providing a robust cluster size for subsequent analyses. Additionally, *deepSplit=2* is set, representing a median sensitivity level for cluster separation, which helps in discerning distinct gene modules.

***Step 3: Summarize the expression profiles in a module – define eigengenes***

For each identified module, we computed the module eigengene, defined as the first principal component of the gene expression profiles within the module. This eigengene effectively summarizes the module's overall expression pattern. To refine the module granularity and improve the interpretability of the results, we performed hierarchical clustering on the module eigengenes using dissimilarity measures (1 - correlation). We applied a cut height of 0.1 to this dendrogram to identify and merge similar modules. The clustering and merging process is illustrated in **Supplemental Figure 7**, where the hierarchical clustering dendrograms are shown both before and after module merging. This approach not only categorizes genes into functionally coherent groups but also facilitates the examination of gene networks in relation to phenotypic traits.

*Step 4: Relate modules to external traits*

We utilized the human organ RNA-Seq time-series of the development of seven major organs (ArrayExpress Accession Number E-MTAB-6814)[1]. The dataset covers 23 developmental stages across 7 organs (n=313: brain/forebrain 55, hindbrain/cerebellum 59, heart 50, kidney 40, liver 50, ovary 18, and testis 41), starting at 4 weeks post-conception until 58-63 years of age. The sample metadata can be downloaded from ArrayExpress - the MAGE-TAB file is `E-MTAB-6814.sdrf.txt` (see https://www.ebi.ac.uk/biostudies/arrayexpress/studies/E-MTAB-6814). This data was used to create a binary matrix representing the seven organs in the dataset, with rows corresponding to samples and columns to tissues. We calculated the correlation between module eigengenes and the binary tissue matrix to identify modules that are specifically expressed in certain tissues. Modules with high correlations to particular tissues, such as the heart, were highlighted as tissue-specific using several data visualizations.

A heatmap was generated to visualize the correlation between module eigengenes and different tissues (**Figure 3**). The color intensity indicated the strength and direction of the correlations, with specific focus on tissue-specific modules. This visualization facilitated the identification of modules that may be critical for the development and function of particular tissues.

We visualized the relationships between module eigengenes across different organs using a dendrogram (**Supplemental Figure 8**) and an adjacency heatmap (**Supplemental Figure 9**). The dendrogram showed the hierarchical clustering of eigengenes, indicating similarity in expression

---

[1] Cardoso-Moreira M, Halbert J, Valloton D, Velten B, Chen C, Shao Y, Liechti A, Ascenção K, Rummel C, Ovchinnikova S, Mazin PV, Xenarios I, Harshman K, Mort M, Cooper DN, Sandi C, Soares MJ, Ferreira PG, Afonso S, Carneiro M, Turner JMA, VandeBerg JL, Fallahshahroudi A, Jensen P, Behr R, Lisgo S, Lindsay S, Khaitovich P, Huber W, Baker J, Anders S, Zhang YE, Kaessmann H. **Gene expression across mammalian organ development.** *Nature*. 2019 Jul;571(7766):505-509. doi: 10.1038/s41586-019-1338-5. PMID: 31243369; PMCID: PMC6658352.

patterns across tissues, while the adjacency heatmap displayed the pairwise correlations between module eigengenes. These visualizations helped identify modules with related expression profiles, potentially sharing biological functions or co-regulated gene sets.

We extended the analysis by correlating module eigengenes with the developmental stages of the heart. Treating the developmental stages as a continuous time series is a powerful approach to modeling the relationship between module eigengenes and the continuous variable developmental age, representing the progression from conception to adulthood. This approach will capture trends and correlations over time, rather than just between categorical stages (i.e., the organs). The heart samples' age post-conception in weeks was calculated using the sample metadata. In humans, the average number of days from conception to birth is approximately 266, equivalent to about 38 weeks. Using 38 weeks for gestation, the subject's age in weeks was calculated for all development stages from conception to adulthood. Correlation of the module eigengenes with heart sample age was performed using the WGCNA::cor function.

For visualization and clustering, the sample's age in weeks was converted into a discrete variable of developmental stage as follows:

- **Early Development (4–10 weeks post-conception):** This stage involves the initial formation of the heart, including the heart tube, the onset of the heartbeat, and the development of primitive atria, ventricles, and outflow tracts.
- **Mid-Development (11–20 weeks post-conception)**: The heart undergoes structural refinement during this period, with the septation of chambers, development of the conduction system, and initiation of blood circulation.
- **Late Fetal Development (21–40 weeks post-conception):** The heart continues to mature in preparation for birth, with muscle maturation, further development of the conduction

system, and adaptation for postnatal circulation. Note: No samples were available for this stage.

- **Perinatal Period (41–52 weeks post-conception):** This stage encompasses the transition around birth, including the closure of fetal circulatory shunts and the final adaptation of the heart structure and function to adult-like circulation patterns.
- **Postnatal Development (1–5 years):** After birth, the heart grows and adapts, with continued myocardial development and stabilization of adult-like function and structure.
- **Mature Heart (>5 years):** The heart has reached full structural and functional maturity, maintaining function and adapting to physical and metabolic demands throughout adulthood.

As with the organs, developmental stage traits were encoded as a binary matrix, each column represented a specific stage, and each row corresponded to a sample. We calculated the correlation between module eigengenes and the external traits using Pearson correlation. We visualized the relationships between module eigengenes across different developmental stages in the heart using a heatmap (**Supplemental Figure 13**), a dendrogram (**Supplemental Figure 14**), and an adjacency heatmap (**Supplemental Figure 15**).

### *Gene Type Enrichment Analysis in WGCNA Modules*

We assessed the enrichment or depletion of lncRNAs, protein-coding genes, and CHD genes within each WGCNA module using hypergeometric and Fisher's exact tests. The approach was as follows:

1. **Data Aggregation:** First, the gene counts were summarized within each module, categorizing them by gene type (lncRNA, protein-coding gene, CHD gene). The total number of genes, lncRNAs, protein-coding genes, and CHD genes across all modules is calculated to provide a reference for the enrichment analysis.

2. **Hypergeometric Test:** The hypergeometric test is performed to calculate the probability of observing the given number of each gene type (lncRNA, protein-coding gene, CHD gene) within each module, relative to the total number of that gene type across all modules. This test helps identify whether the observed counts of these gene types in each module are higher or lower than expected by chance.
3. **Fisher's Exact Test:** Fisher's exact test is applied to each module to determine whether the counts of lncRNAs, protein-coding genes, and CHD genes are significantly enriched or depleted compared to what would be expected by random distribution. The results include both p-values and odds ratios. The p-values from Fisher's exact test are adjusted for multiple testing using the Benjamini-Hochberg method, controlling the false discovery rate.
4. **Enrichment/Depletion Classification:** Based on the adjusted p-values and odds ratios from Fisher's exact test, each gene type within each module is classified as "Enriched," "Depleted," or "Not Significant."

***Module-specific Gene Ontology term enrichment analysis.***

We applied the guilt-by-association method to infer the functions of lncRNAs included in modules by analyzing their association with correlated protein-coding genes important in heart development or causally associated with CHDs. This approach assumes that lncRNAs may share functional characteristics with closely related protein-coding genes in the same module. For the functional enrichment analysis, we exclusively used protein-coding genes from the target modules because Gene Ontology (**GO**) terms typically do not cover lncRNA genes.

Using the *clusterProfiler* tool (v4.12.6), we applied the *enrichGO* function to determine enriched GO terms across three categories: biological processes (**BP**), molecular function (**MF**), and cellular component (**CC**). We adjusted the p-values ($\mathbf{p_{adj}}$) using the Benjamini-Hochberg method, setting a significance threshold of p-value < 0.01 and q-value < 0.05.

We further refined the GO terms by calculating semantic similarity using the Wang method, which integrates the hierarchical structure of GO terms to provide biologically meaningful similarity scores. For each ontology (BP, MF, CC), the semantic similarity between GO terms was calculated using the termSim() function from the GOSemSim package. Information Content (IC) was added to each enriched term using godata(), based on ENSEMBL gene annotations.

To reduce redundant GO terms, we applied hierarchical clustering on the semantic similarity matrix. The similarity matrix was first transformed into a distance matrix by subtracting the similarity scores from 1. We removed rows and columns where all values were NA, and the resulting distance matrix was used to perform clustering using the average linkage method (hclust). The dendrograms for each ontology were cut at a height of 0.3 to form clusters of similar GO terms. Within each cluster, we selected representative GO terms based on the lowest adjusted p-value (p.adjust). The resulting representative terms from the three GO categories were combined into a final set of non-redundant GO terms. Finally, the adjusted p-values (p.adjust) for the representative terms were transformed into -log10 scale for easier interpretation. The resulting terms were ranked by their significance (adjusted p-value), and the most significant terms were retained for further analysis.

***Identification of Hub Genes***

We focused on identifying hub genes within specific modules by examining the relationship between Module Membership (**MM**) and Gene Significance (**GS**). The process involved the following key steps:

1. **Module Membership (MM)**: We calculated the module eigengene (the first principal component of the gene expression profiles) for each identified module. MM values were then

computed for each gene, reflecting how strongly each gene is associated with its module eigengene.

2. **Gene Significance (GS)**: We calculated the correlation between each gene's expression and the heart trait to determine GS values, representing how strongly each gene is associated with the trait.
3. **Visualizing MM vs. GS**: Scatter plots were generated to visualize the relationship between MM and GS for each module, helping to identify hub genes—those with high MM and GS values. These hub genes are key drivers of the biological processes represented by the modules.

**GINI Coefficient**

VanOudenhove *et al.* (2020)[2] use the **Gini coefficient** to measure gene expression specificity across tissues during human heart development (see online Supplementary File V). The Gini coefficient, traditionally used to measure income inequality, is applied here to assess the tissue-specific expression of genes. In the study the Gini index is used to quantify how uniformly or specifically a gene is expressed across different tissues. Genes with high Gini coefficients (e.g., greater than 0.5) are identified as being more tissue-specific—in this case, specific to the embryonic heart. These genes are enriched for heart-related functions, including heart development and associated processes. For example, they identified 347 genes with a Gini coefficient greater than 0.5 that had heart-specific expression, which included well-known heart development genes like *TBX5*, *IRX4*, and *HAND1*.

---

[2] VanOudenhove J, Yankee TN, Wilderman A, Cotney J. **Epigenomic and Transcriptomic Dynamics During Human Heart Organogenesis.** *Circ Res.* 2020 Oct 9;127(9):e184-e209. doi: 10.1161/CIRCRESAHA.120.316704. Epub 2020 Aug 9. PMID: 32772801; PMCID: PMC7554226.

The Gini coefficient helps identify genes likely important in heart development by highlighting those with expression patterns specific to the heart during embryogenesis. To compare to the hub genes identified in our analysis, the LncBook identifiers were mapped to Ensembl gene accessions using a conversion file provided by the LncBook developers (LncBook_id_conversion.csv). For conversion of smaller batches of lncRNAs, the online LncBook batch conversion tool can be used.

Hub genes in our dataset were then annotated with the VanOudenhove *et al.* Gini index. For the 18 lncRNAs in the hub genes of the magenta module (ME03), 7 of the 9 that could be mapped to an Ensembl accession number have a Gini coefficient. Of the 8 known CHD genes in the module, all had Ensembl accessions and Gini coefficients. Ten genes (five CHD genes and five lncRNA genes) with maximal expression in the heart had a Gini coefficient, 9 of which were significant (> 0.5).
