## Supplementary Figures for "Identification of Long Non-coding RNA Candidate Disease Genes Associated with Clinically Reported CNVs in Congenital Heart Disease"

† Cytogenomics of Cardiovascular Malformations (CCVM) Consortium – see **Supplemental Methods** for the complete list of Contributing Authors.

**Short title**: CNVs impacting lncRNAs in congenital heart disease

***Corresponding Author:** Peter White, Ph.D., Battelle Endowed Chair for Quantitative and Computational Medicine, The Office of Data Sciences, The Abigail Wexner Research Institute at Nationwide Children's Hospital, 575 Children's Crossroad, Columbus, OH 43215. USA.

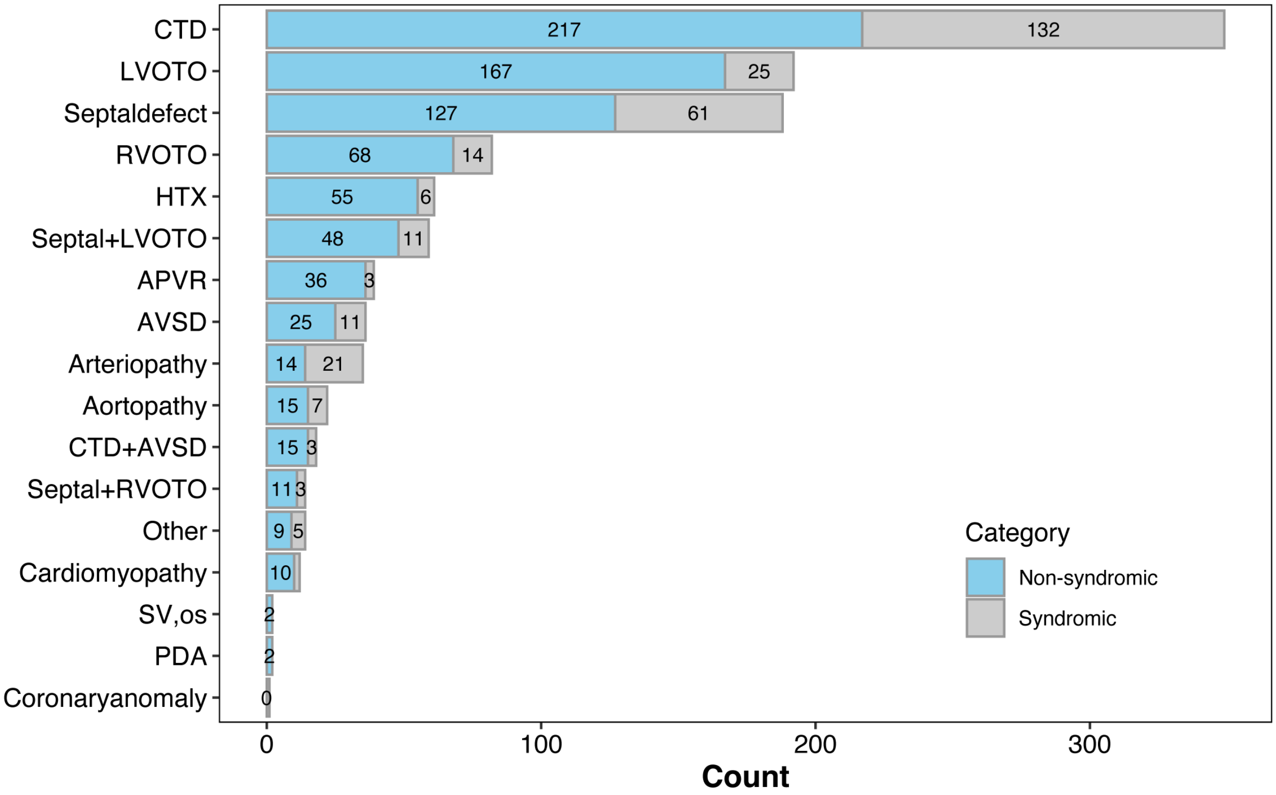


**Supplemental Figure 1: Counts of Syndromic and Non-syndromic Patients by Diagnosis.** This bar plot visualizes the distribution of syndromic and non-syndromic patients across various CHD diagnoses. Each bar represents the total number of patients within a specific diagnosis, with the count of non-syndromic patients shown in sky blue and syndromic patients in grey. The bars are stacked, allowing for easy comparison of the relative proportions of syndromic and non-syndromic cases within each diagnosis. The diagnoses are listed on the y-axis, while the total patient count is shown on the x-axis. This visualization highlights the variation in syndromic and non-syndromic occurrences across different CHD categories in the study cohort.

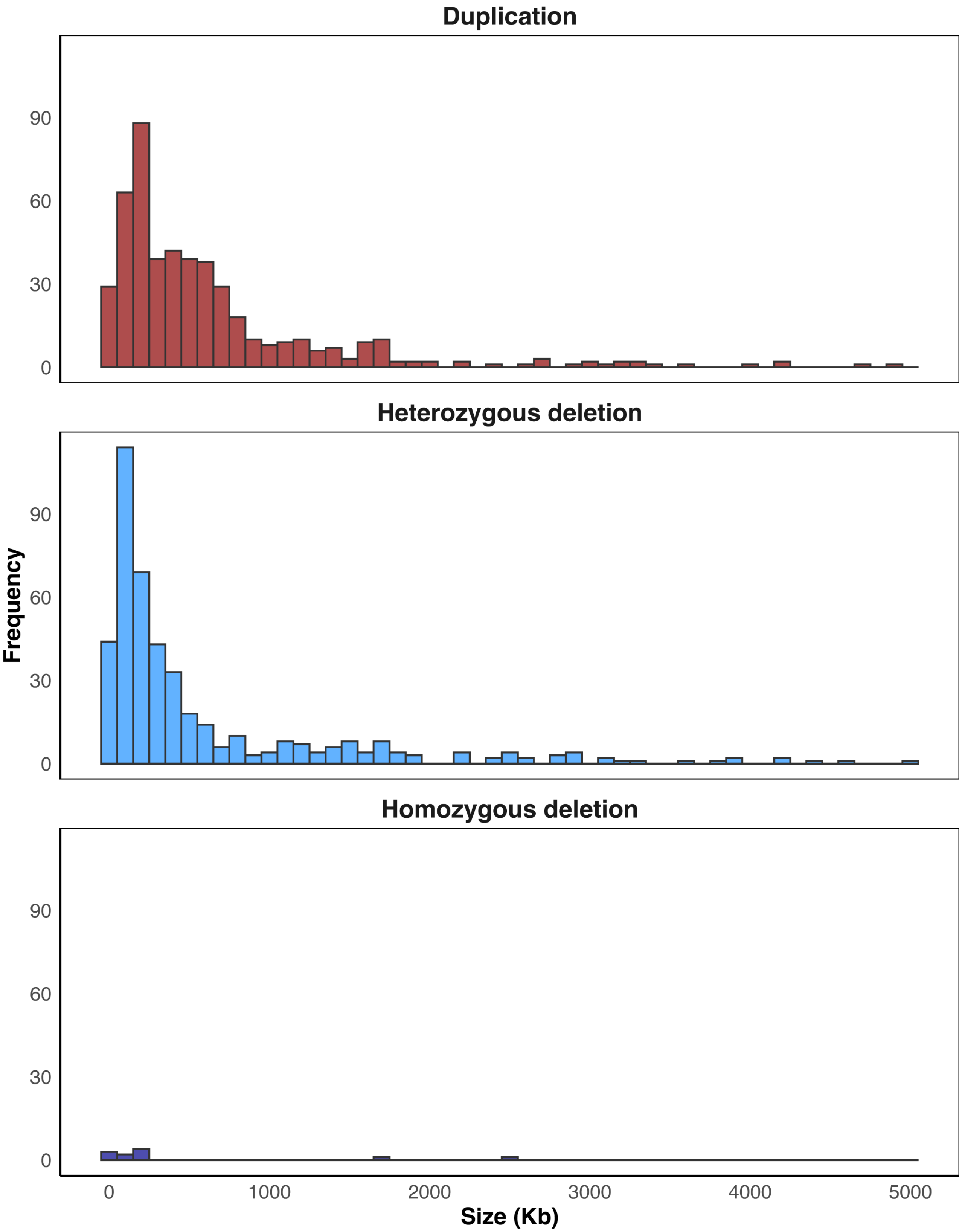


**Supplemental Figure 2: CNV Size Distributions Across Isolated CHD Patients.** This histogram illustrates the distribution of CNV sizes within the isolated CHD cohort. CNV sizes are measured in kilobases (Kb) and are displayed on the x-axis, while the frequency of each size range is shown on the y-axis. The histogram is faceted by CNV value, with each panel representing a different CNV type: duplication (red), heterozygous deletion (blue), and homozygous deletion (purple). This plot provides a detailed view of the variation in CNV sizes across the cohort, highlighting the most common size ranges within each CNV category.

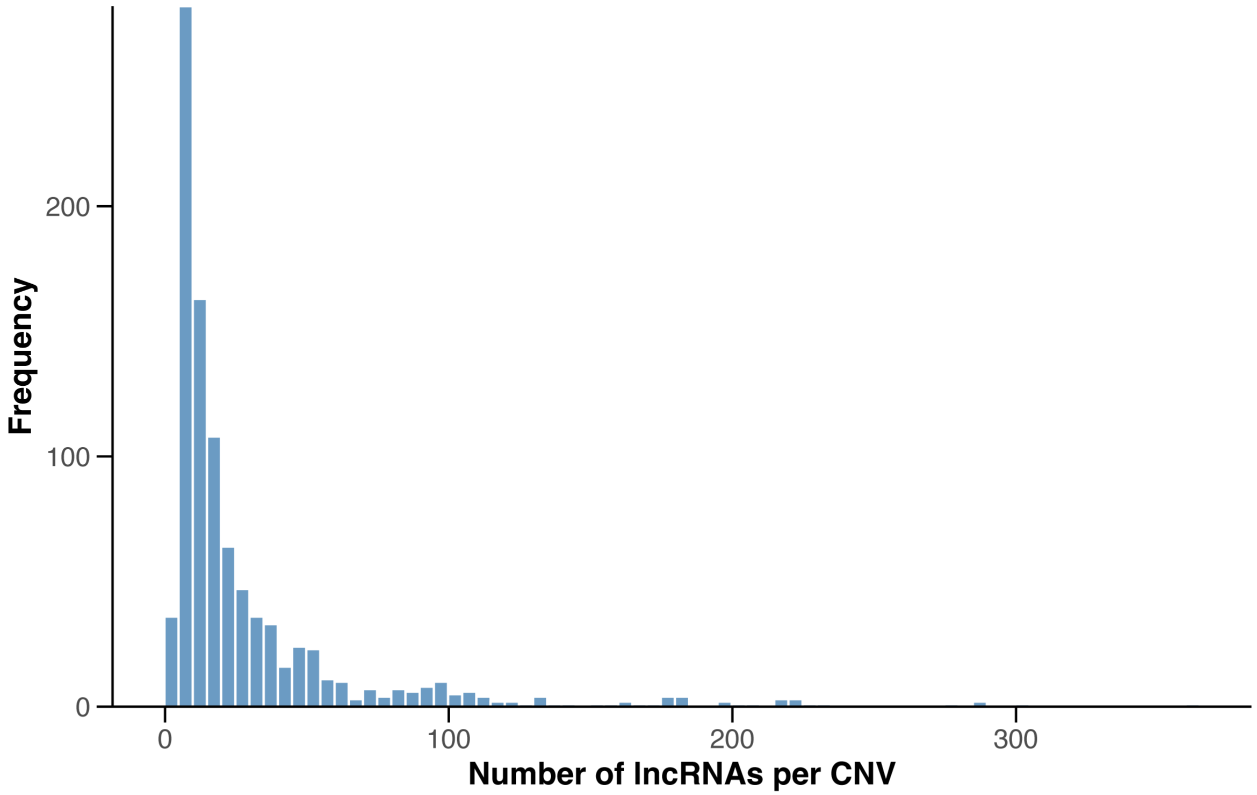


**Supplemental Figure 3: Distribution of lncRNA Counts Within CNVs.** This histogram illustrates the distribution of CNV-lncRNAs in isolated CHD patients within the Cytogenomics of Cardiovascular Malformations (CCVM) Consortium cohort. CNVs are grouped into bins based on the number of overlapping lncRNAs, with zero counts handled separately. The x-axis represents the number of lncRNAs per CNV, while the y-axis shows the frequency of CNVs in each bin. The plot highlights the prevalence of CNVs with varying levels of lncRNA involvement, providing insights into the genomic regions most frequently associated with lncRNAs.

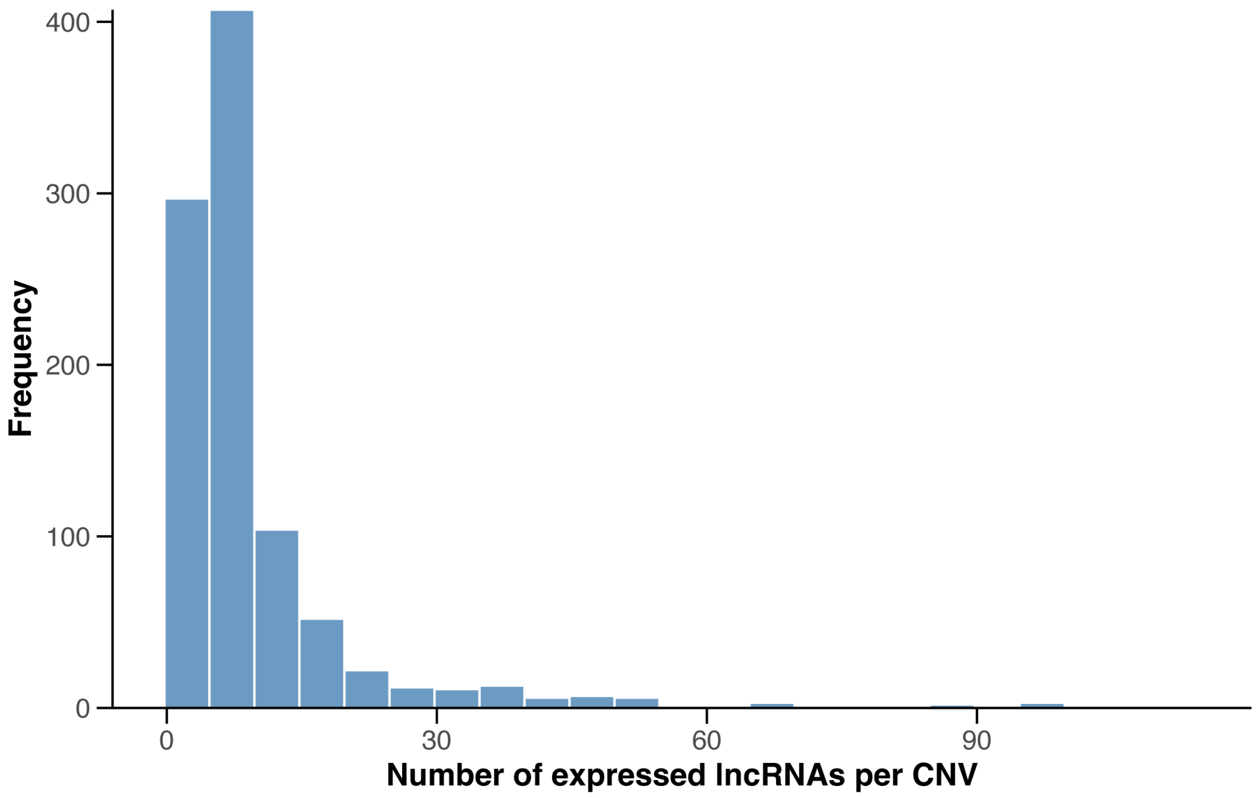


**Supplemental Figure 4: Distribution of Expressed lncRNA Counts Within CNVs.** This histogram illustrates the distribution of the number of expressed CNV-lncRNAs in the CCVM study cohort. Expression was determined by a TPM value >=1 in any of the 50 heart samples from the developmental time series RNA-seq data. CNVs are grouped into bins based on the number of overlapping lncRNAs, with zero counts handled separately. The x-axis represents the number of lncRNAs per CNV, while the y-axis shows the frequency of CNVs in each bin. The plot highlights the prevalence of CNVs with varying levels of lncRNA involvement.

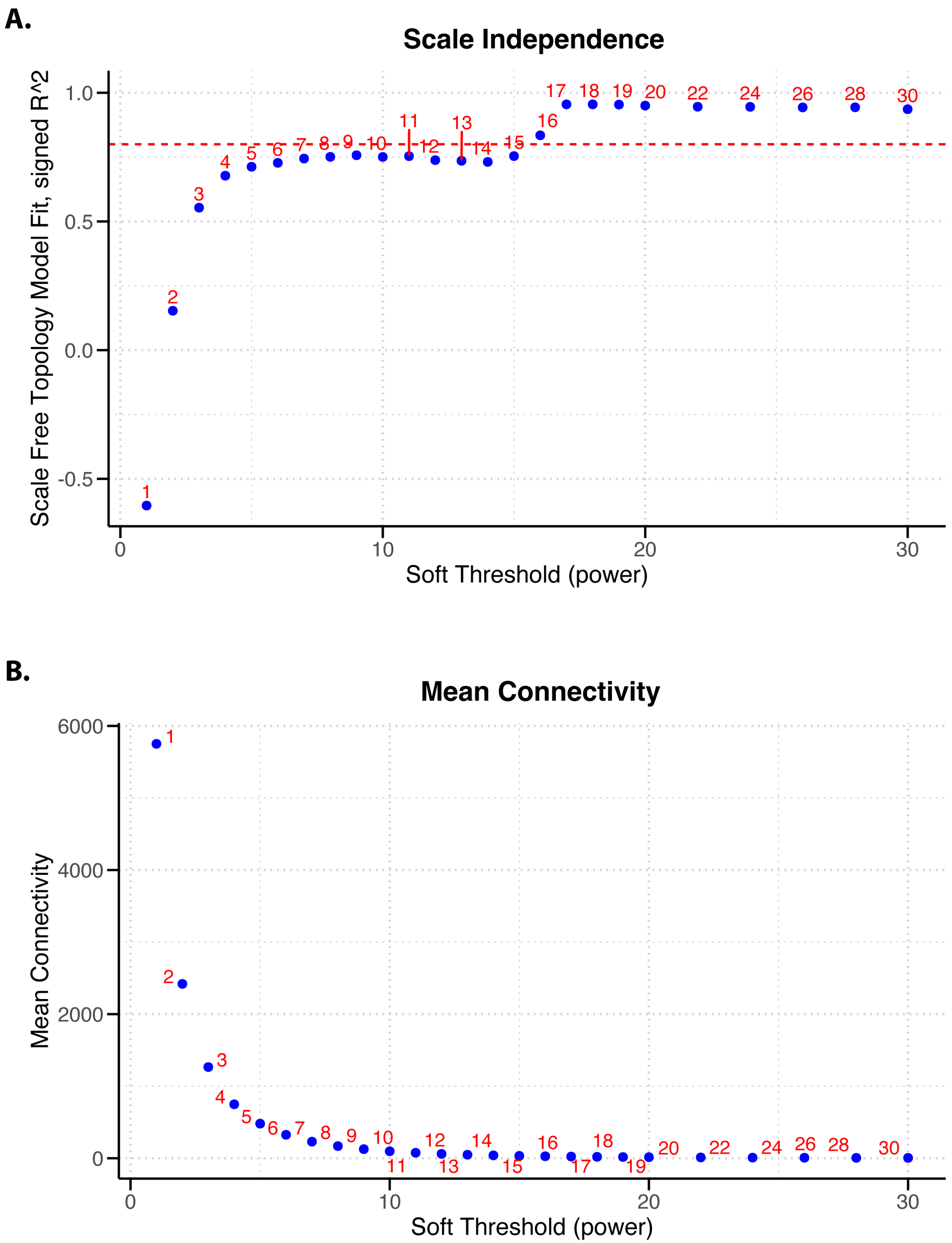


**Supplemental Figure 5: Selection of Soft-Thresholding Power for WGCNA Network Construction.** (A) Scale Independence Plot: This plot shows the scale-free topology model fit ($R^2$) as a function of the soft-thresholding power. The red dashed line indicates the threshold of $R^2 = 0.80$, which is typically considered sufficient for achieving a scale-free network. Each point represents the $R^2$ value for a corresponding power, with the selected power highlighted. (B) Mean Connectivity Plot: This plot displays the mean connectivity of the network as a function of the soft-thresholding power. Mean connectivity represents the average number of connections each node has to other nodes. The selected power is chosen to balance a high $R^2$ value with reasonable network connectivity, ensuring that the network is both scale-free and sufficiently connected.

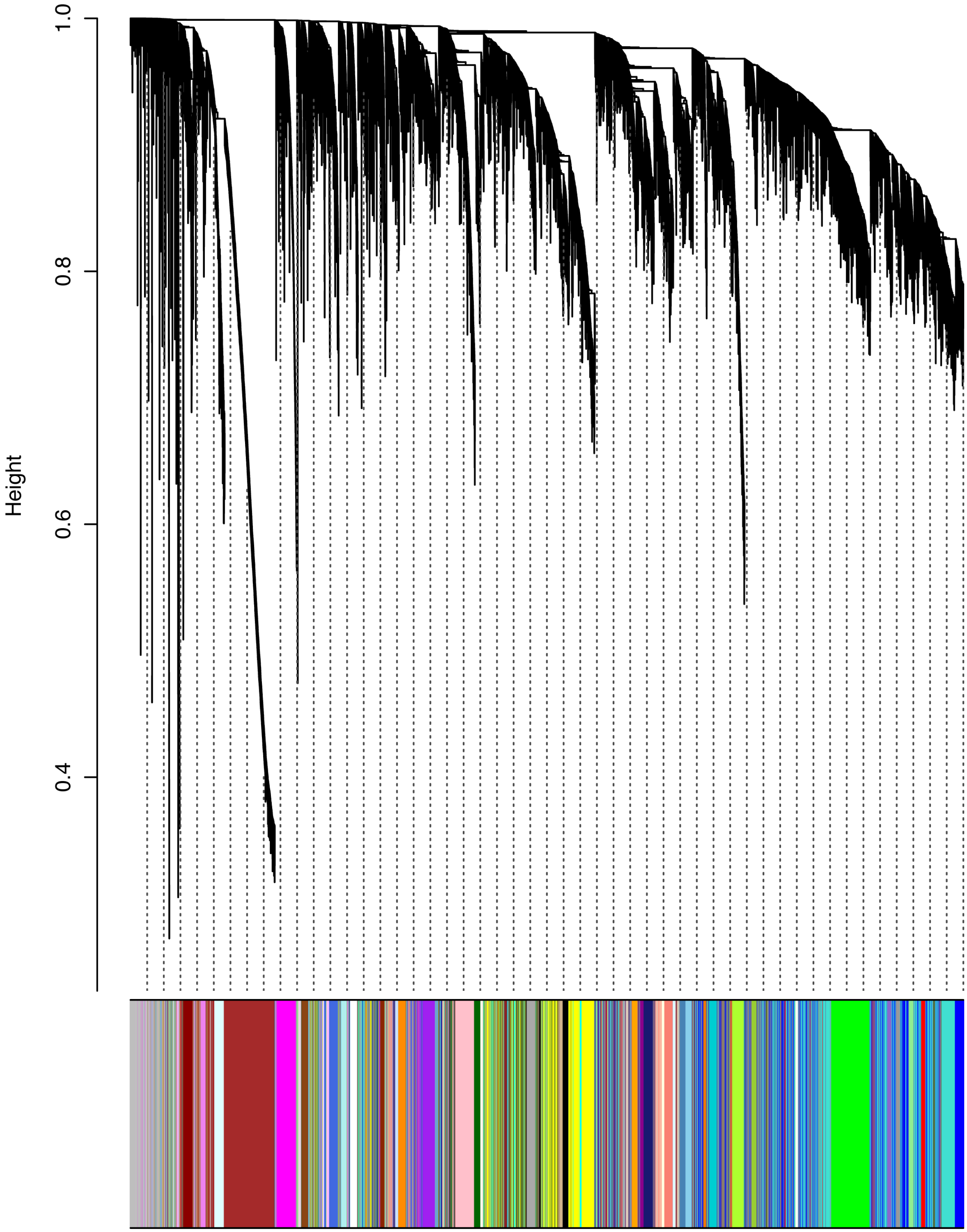


**Supplemental Figure 6: Gene Dendrogram and Module Assignment Based on Dynamic Tree Cut.** The dendrogram represents the hierarchical clustering of genes based on the Topological Overlap Matrix (TOM)-based dissimilarity measure. Each branch of the dendrogram corresponds to a group of genes with similar expression patterns. Below the dendrogram, the dynamic tree-cut method has been applied to identify distinct gene modules represented by different colors. Each color indicates a module, a cluster of co-expressed genes, and potentially functionally related genes. The vertical guidelines indicate the modules detected by the dynamic tree-cut method, with the height of the dendrogram reflecting the dissimilarity between genes. This visualization helps to identify and interpret the modular organization of the gene co-expression network.

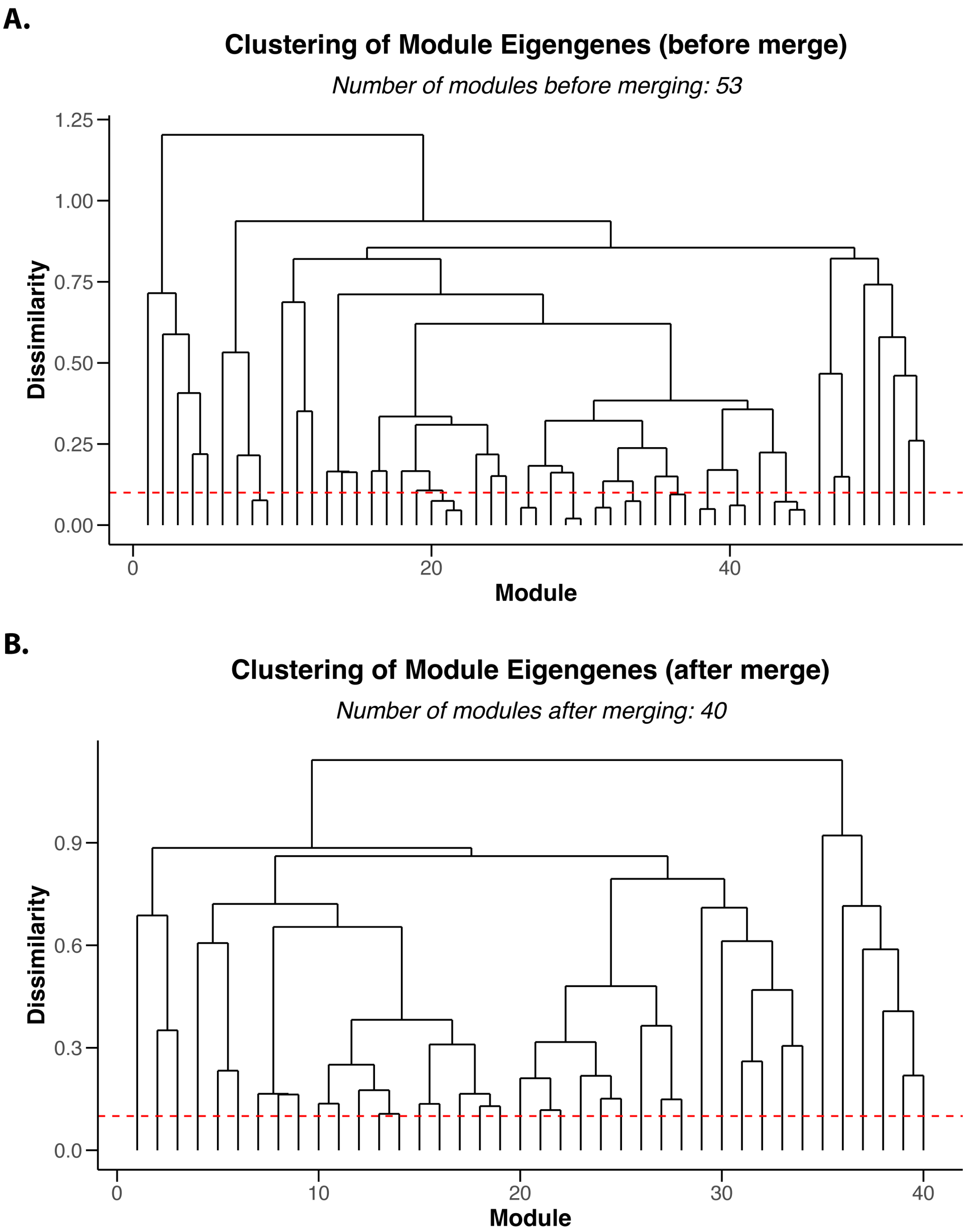


**Supplemental Figure 7: Hierarchical Clustering of Module Eigengenes Before and After Merging.** (A) Clustering of Module Eigengenes Before Merging: The dendrogram shows the hierarchical clustering of module eigengenes based on their dissimilarity, calculated as 1 - correlation. The red dashed line represents the cut height (0.1) used to identify similar modules to be merged. Before merging, there are 53 distinct modules. (B) Clustering of Module Eigengenes After Merging: The dendrogram shows the hierarchical clustering of the module eigengenes after merging similar modules. After merging, the number of distinct modules is reduced to 40. This demonstrates how closely related modules have been combined to simplify the network while preserving key relationships.

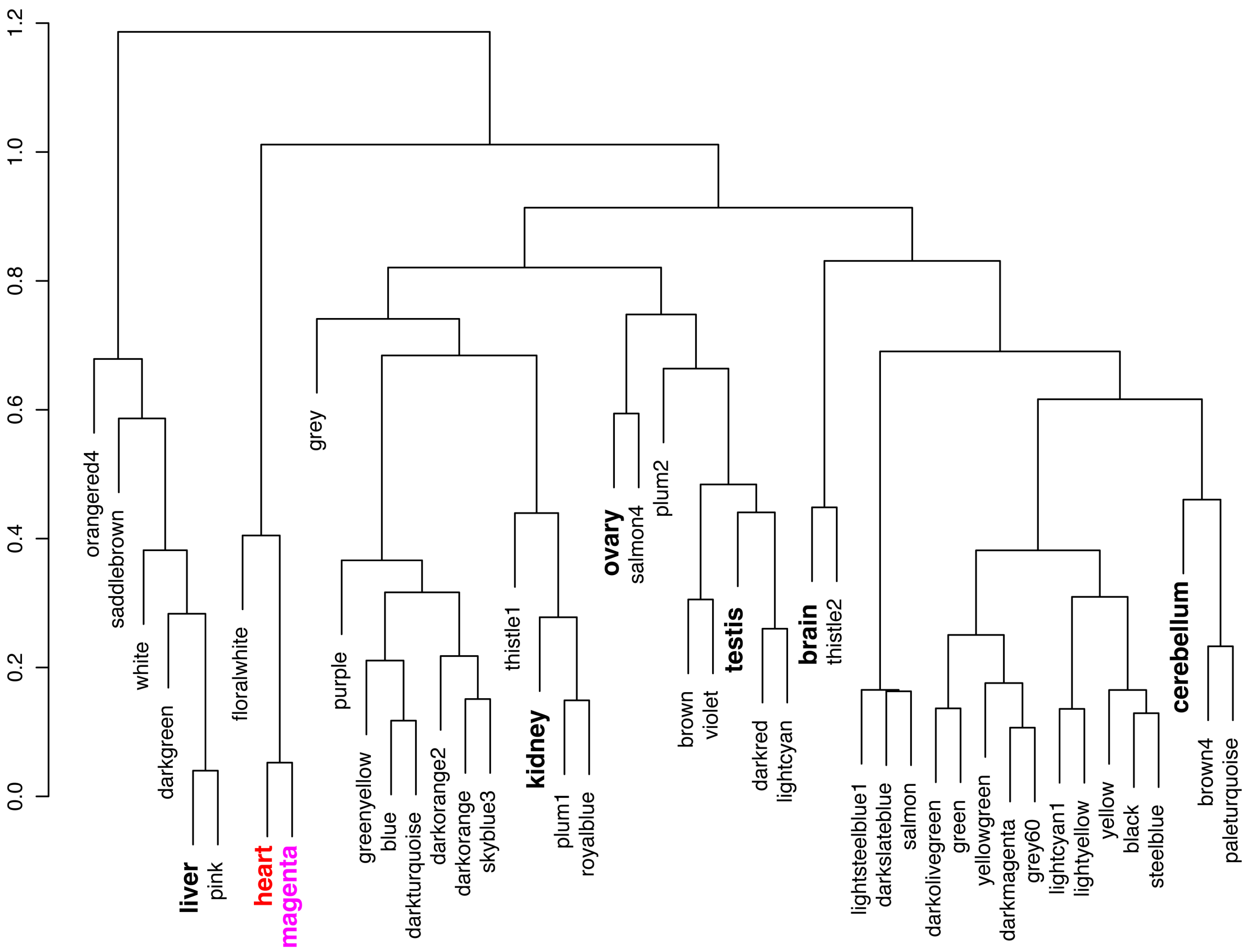


**Supplemental Figure 8: Module Eigengene Dendrogram for Organ-Specific Modules.** This dendrogram illustrates the hierarchical clustering of module eigengenes across different organs. Each branch represents a module eigengene, and the clustering reflects the similarity in expression patterns between the modules. Modules that are closely related are grouped together on the dendrogram, indicating that they have similar expression profiles across the organs. This visualization helps to identify which modules are most closely associated with each other, and to which organs (bold), providing insights into potential shared biological functions or co-regulated gene sets across different tissues. The clustering of the magenta module (ME03) and the heart is highlighted by color.

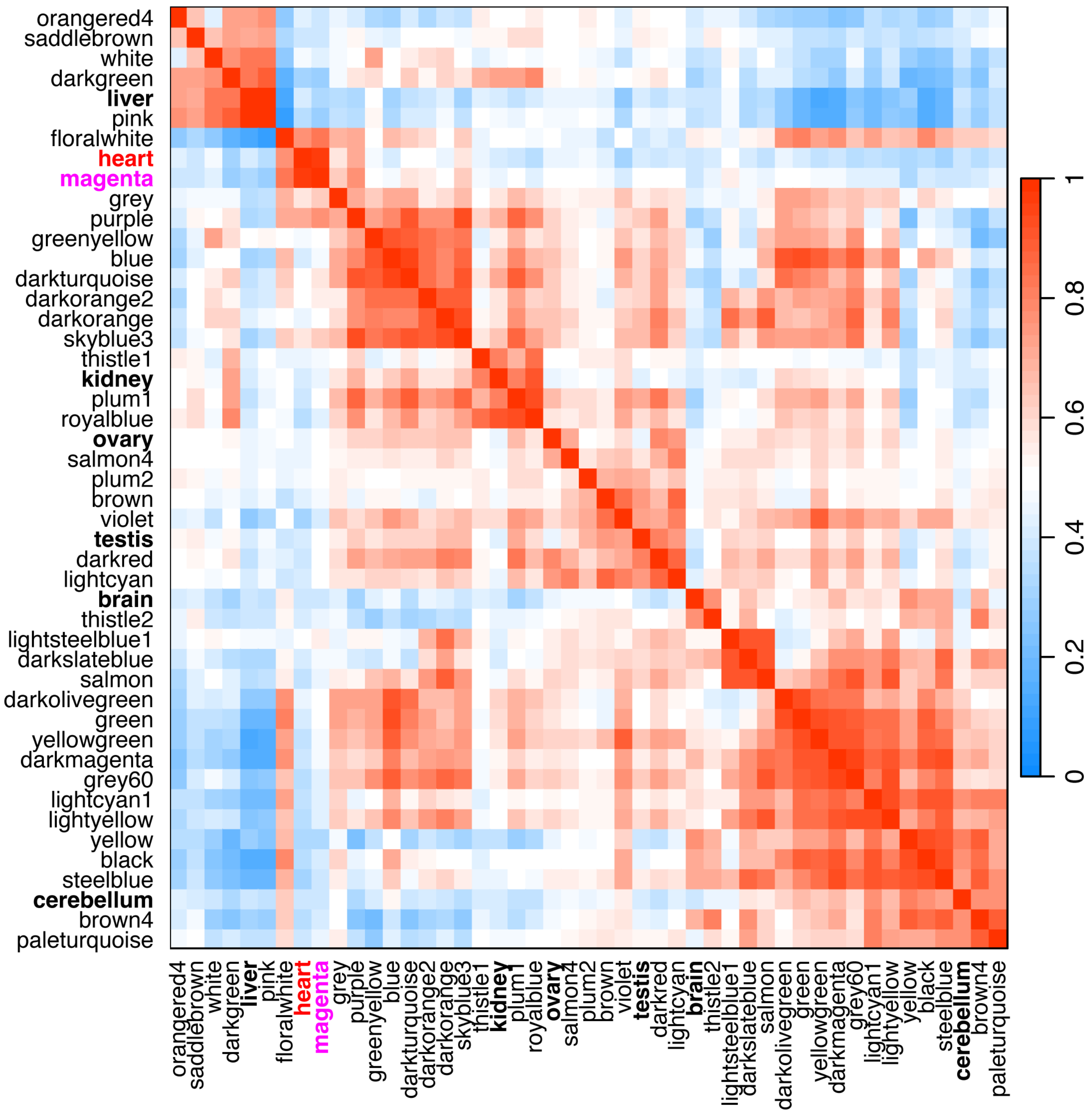


**Supplemental Figure 9: Module Eigengene Adjacency Heatmap for Organ-Specific Modules.** This heatmap displays the pairwise correlations between module eigengenes across different organs, with color intensity reflecting the strength and direction of the correlation. Positive correlations are shown in red, indicating that the modules have similar expression patterns, while negative correlations are shown in blue, indicating opposing expression patterns. The heatmap allows for a visual comparison of how closely related the modules are regarding their expression across various organs, highlighting potential tissue-specific modules or those co-expressed in multiple tissues.

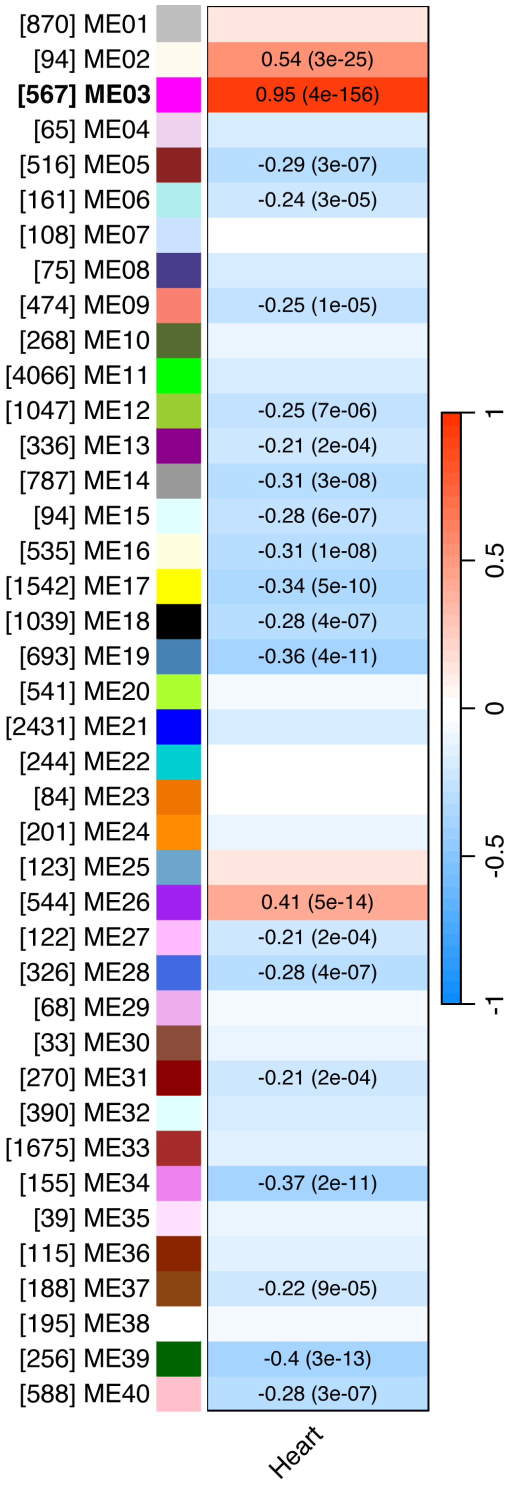


**Supplemental Figure 10: Correlation Between Module Eigengenes and Heart Trait.** This heatmap visualizes the correlation between each module eigengene and the heart trait, with the color intensity reflecting both the strength and direction of the correlation. Each row represents a module eigengene, and the single column corresponds to the heart trait. The color labels on the y-axis indicate the module colors, providing a clear visual connection to the modules identified in earlier steps. The values within each cell display the correlation coefficient (R) and the corresponding p-value, with stronger correlations and more significant p-values highlighted by more intense colors. Modules with high positive correlations (in red) are positively associated with the heart trait, while those with high negative correlations (in blue) are negatively associated. This figure helps to identify which modules are most relevant to heart tissue function, guiding further investigation into the biological significance of these modules.

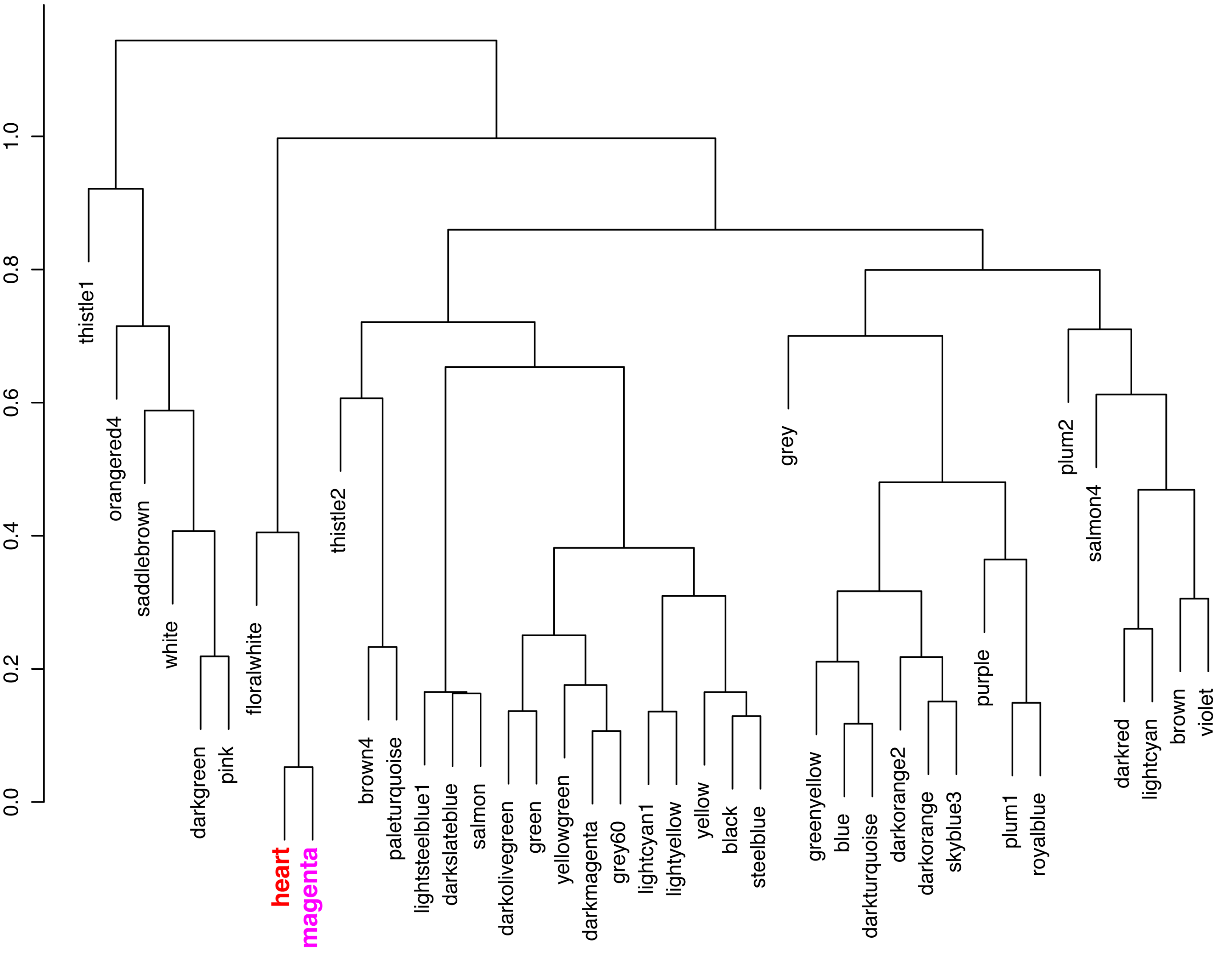


**Supplemental Figure 11: Module Eigengene Dendrogram for Heart-Specific Correlations.** This dendrogram shows the hierarchical clustering of module eigengenes and the heart trait. Each branch in the dendrogram represents a module eigengene or the heart trait, with the clustering reflecting the similarity in expression patterns. Modules closely related in their expression across the heart trait are grouped together, highlighting potential co-regulation or shared functional pathways within heart tissue. This visualization aids in identifying which modules are most strongly associated with the heart trait, offering insights into the molecular networks that may drive heart-specific functions or conditions.

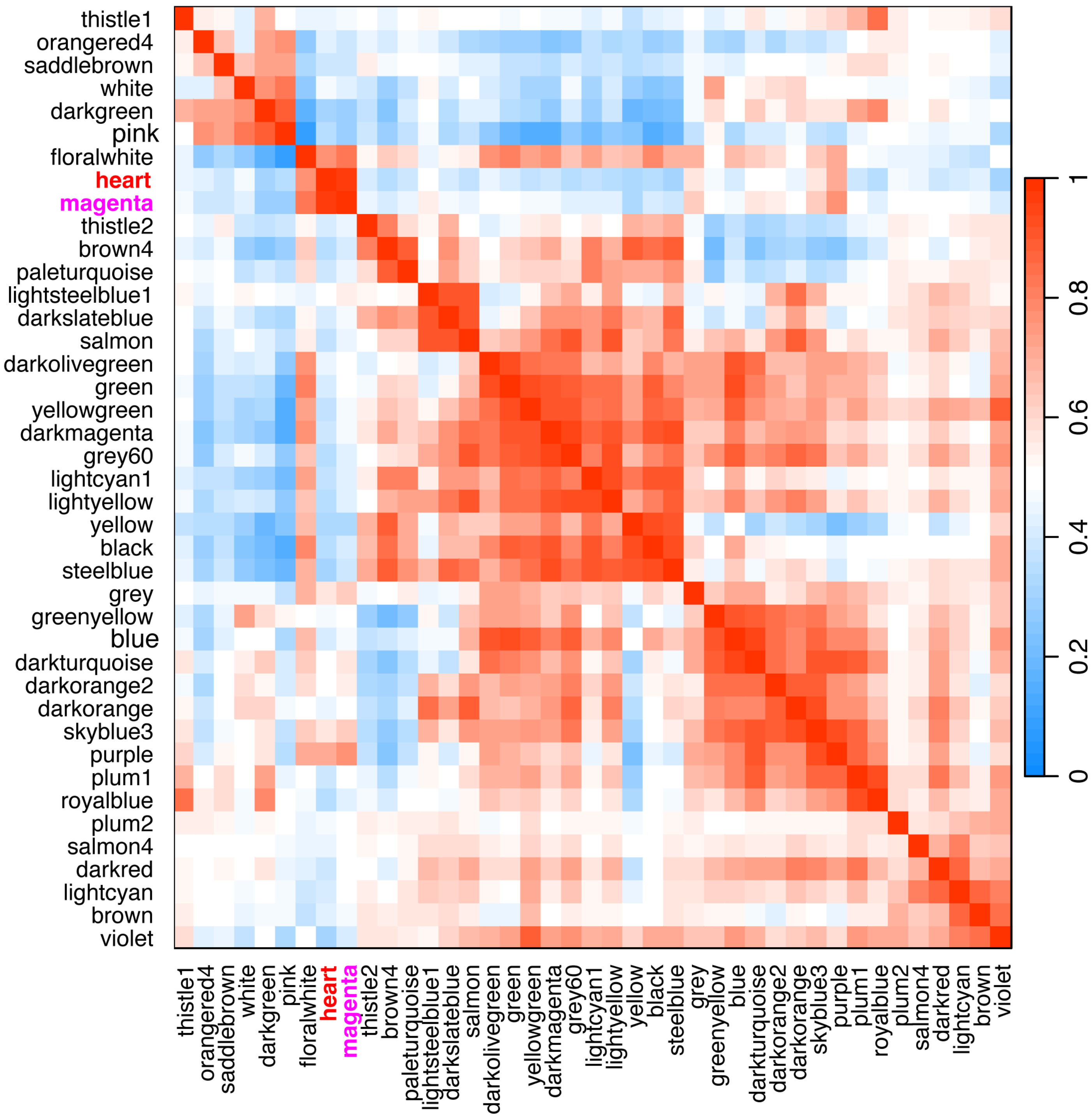


**Supplemental Figure 12: Module Eigengene Adjacency Heatmap for Heart-Specific Correlations.** This heatmap presents the pairwise correlations between module eigengenes and the heart trait, with color intensity representing the strength and direction of the correlations. Positive correlations are depicted in red, indicating modules with expression patterns that are positively associated with the heart trait, while negative correlations are shown in blue. The heatmap provides a detailed view of the relationships between modules in the context of the heart trait, allowing for the identification of modules that may be key regulators or indicators of heart-specific processes. This figure is essential for pinpointing modules with strong heart associations, guiding further analysis of their biological significance in heart tissue.

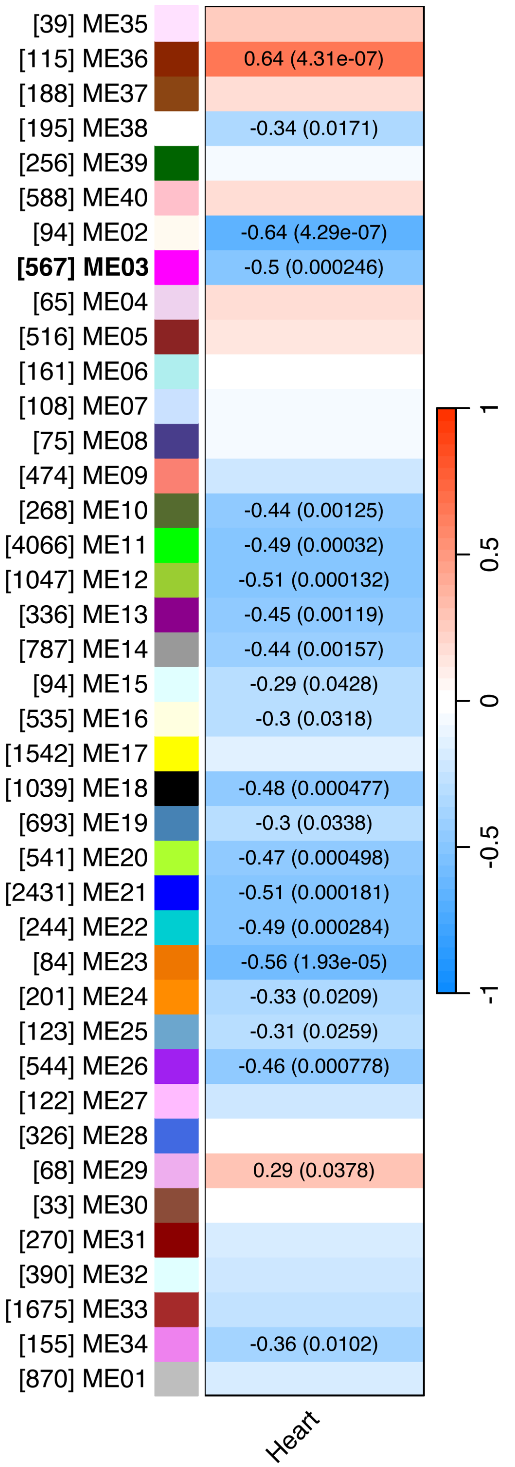


**Supplemental Figure 13: Correlation Between Module Eigengenes and Age in Heart Samples.** This heatmap illustrates the correlation between module eigengenes and age (in weeks) in heart samples, with color intensity reflecting the strength and direction of the correlation. Each row represents a module eigengene, and the single column corresponds to age in heart samples. The y-axis labels include the module name and the number of genes within each module, providing context for the size and identity of each module. Modules with correlation values (|R|) greater than or equal to 0.2 and p-values less than or equal to 0.05 have their correlation coefficients and p-values displayed within the heatmap cells. The color-coded labels on the y-axis correspond to the module colors, helping to link the modules to earlier steps in the analysis visually. Modules with high positive correlations (in red) indicate increasing expression with developmental stage, while negative correlations (in blue) suggest decreasing expression over time. This figure highlights the modules most strongly associated with heart development, identifying key gene networks involved in heart growth and maturation.

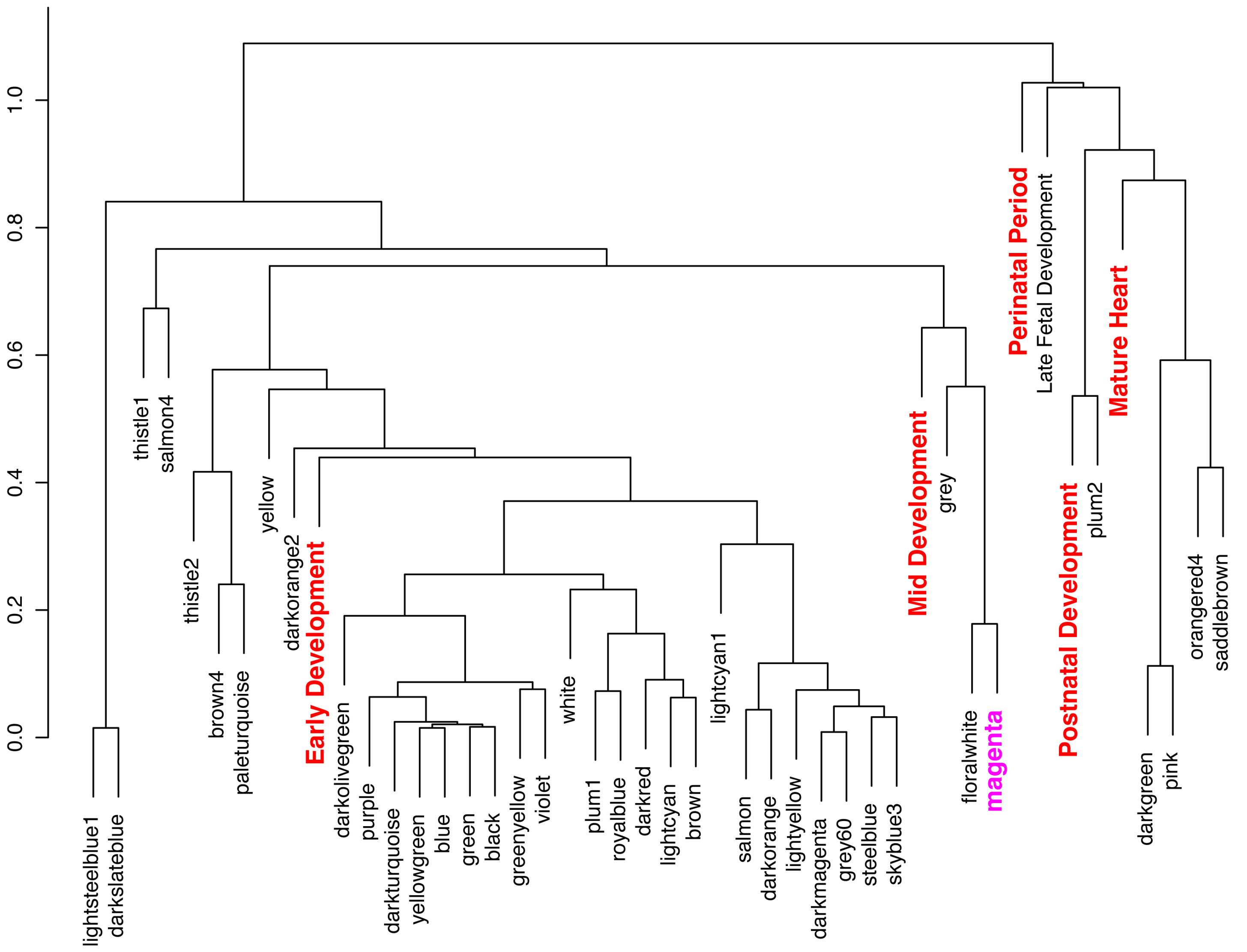


**Supplemental Figure 14: Module Eigengene Dendrogram for Heart Developmental Stages.** This dendrogram illustrates the hierarchical clustering of module eigengenes in heart samples, along with developmental stage groups. Each branch represents a module eigengene or a developmental stage, with the clustering reflecting the similarity in expression patterns across these stages. Modules closely related in their expression during heart development are grouped together, highlighting potential co-regulation or shared functional pathways. This visualization aids in identifying which modules are most strongly associated with specific stages of heart development, offering insights into the molecular networks that drive heart growth and maturation.

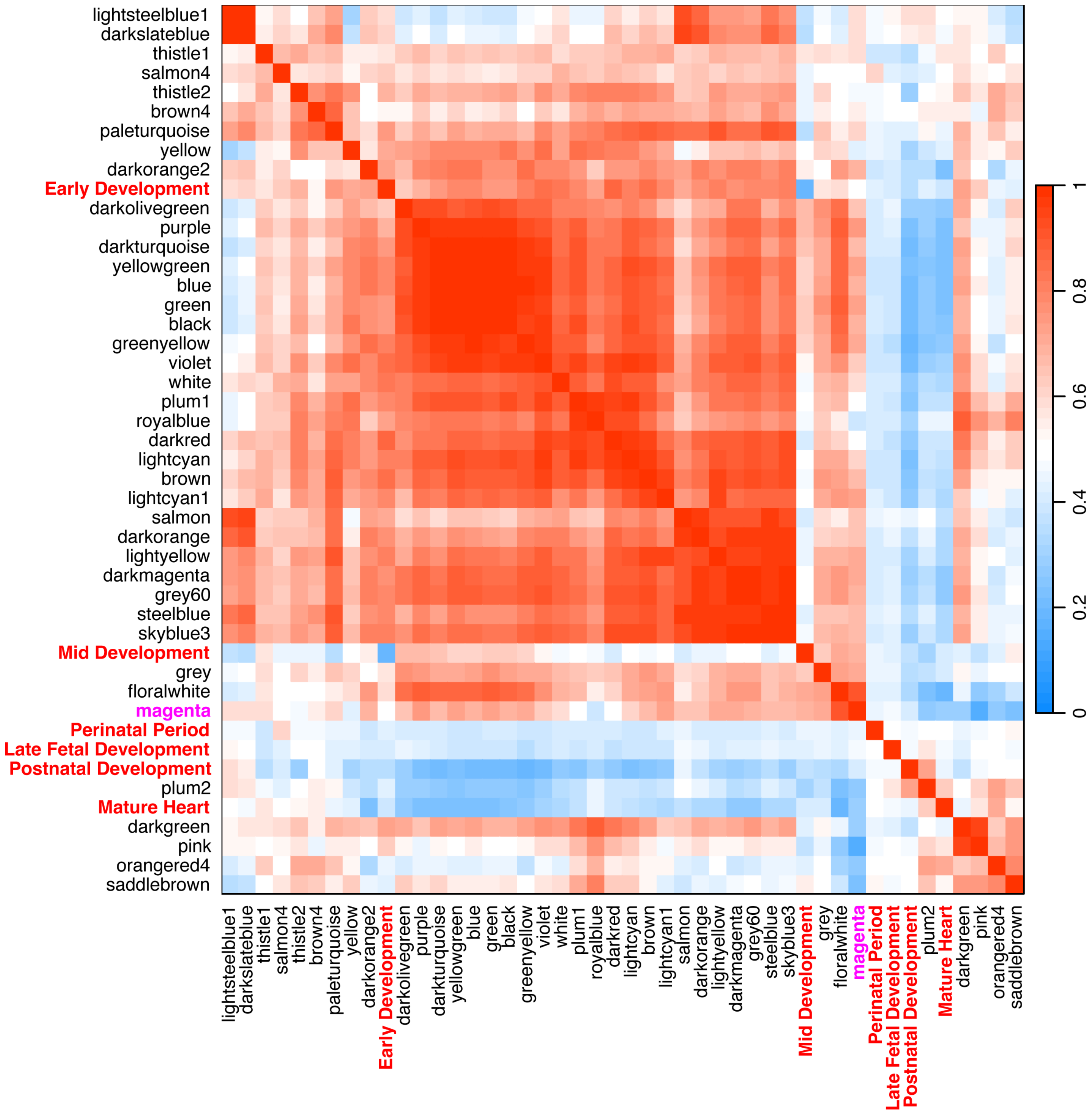


**Supplemental Figure 15: Module Eigengene Adjacency Heatmap for Heart Developmental Stages.** This heatmap presents the pairwise correlations between module eigengenes and developmental stages in heart samples. Color intensity reflects the strength and direction of the correlations, with red indicating positive correlations and blue indicating negative correlations. The heatmap allows for a visual comparison of how closely related the modules are regarding their expression patterns across different stages of heart development. This figure is crucial for pinpointing key modules correlated with specific stages of heart development, guiding further analysis of their biological significance in heart maturation and function.

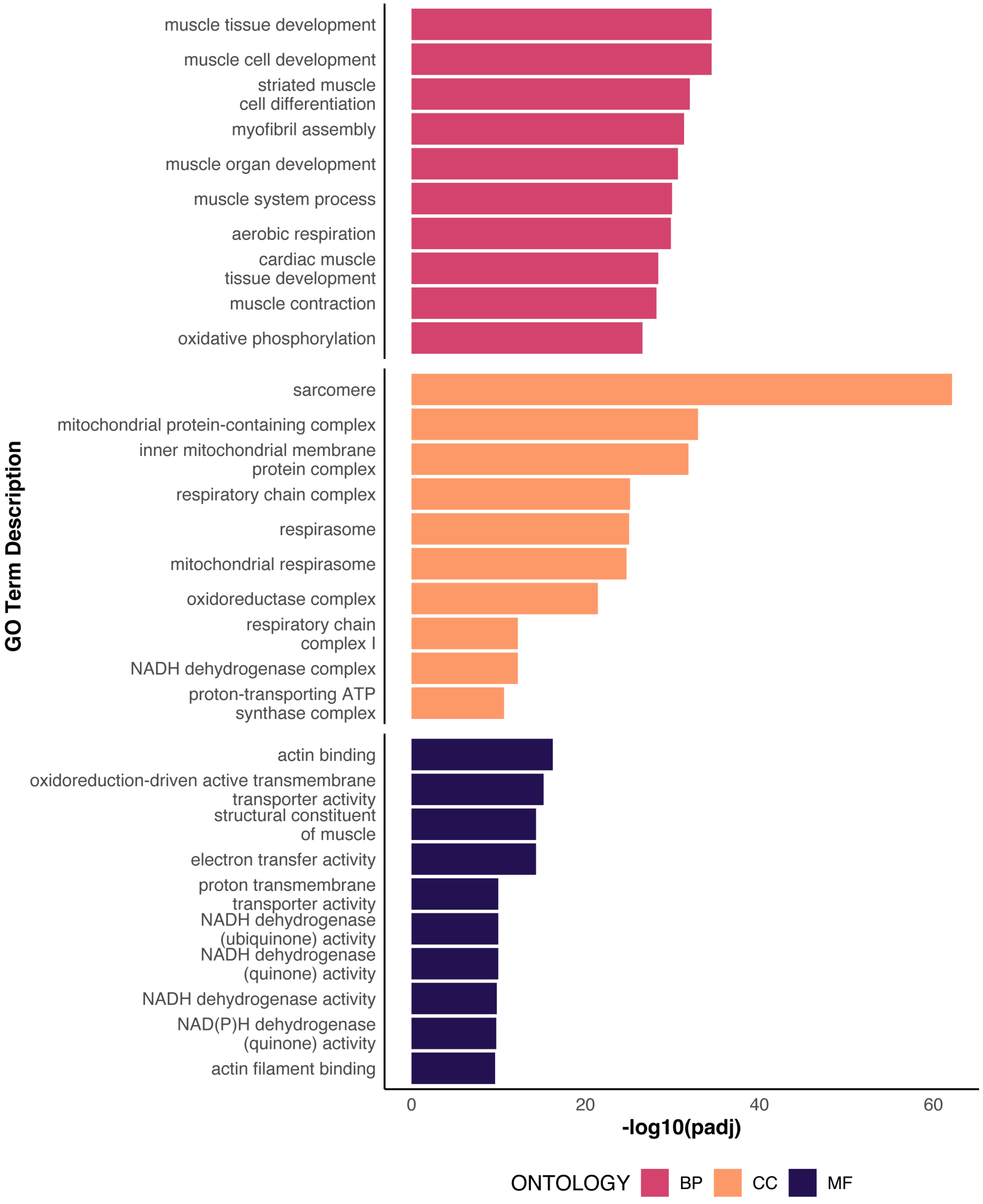


**Supplemental Figure 16: GO Functional Enrichment Analysis for Protein-Coding Genes in the Magenta Module.** This bar chart illustrates the top 10 enriched Gene Ontology (GO) terms across three categories: Biological Process (BP), Molecular Function (MF), and Cellular Component (CC), identified from the protein-coding genes in the magenta module. The GO terms are selected based on the lowest adjusted p-values (Benjamini-Hochberg method), and the bars represent the -log10 transformation of these p-values, providing a clear view of the statistical significance of each term. The terms are color-coded by their GO category. The results highlight key biological processes, molecular functions, and cellular components associated with the magenta module, offering insights into the functional roles of the genes within this module.
